# BioResearchBench: 50 Long Horizon Biology Research Tasks

**DOI:** 10.64898/2026.10.28.762290

**Authors:** Samar Abedrabbo, Alex Pryhoda, Keenan Goodman, Jesse Taavola, Evrhet Milam, Abbey Albright, Reza Jabal, Mi Yang, Cosette Burrese, Justin Tubby, Juan Villagomez, Joe Kim

## Abstract

AI models are used in biology research to study disease mechanisms, discover potential treatments, analyze experimental data, troubleshoot problems, and plan next steps. Useful assistance depends on working with data in various scientific formats, interpreting results, and deciding what to test next. We present BioResearchBench, a benchmark of 50 research questions developed by PhD biologists drawing on their deep research expertise to reflect realistic problems encountered in their fields. Questions were refined through expert review and iterative model testing. The collection spans 10 diverse biology subdomains and includes sequencing data, proteomic measurements, microscopy images, assay readouts, and clinical datasets. We compared recorded outcomes and examined model responses in depth across the 50 tasks on OpenAI GPT-6 Astra, Claude Opus 5.5, Gemini 4 Argon, Meta Muse Wonder, and SpaceXSI Grok 4.7. Success across our benchmark was 18% for Claude Opus 5.5, 13% for Gemini 4 Argon, 11% for Meta Muse Wonder, 4% for GPT-6 Astra, and 4% for Grok 4.7. These results describe a deliberately challenging collection selected through expert review and model testing. Despite low whole task success, the selected responses met most rubric requirements on average: the average fraction of criteria missed ranged from 18% to 32% across models. Across models, the proportion of eligible tasks with at least one category failure was highest in computation and data handling (66.9% failure), followed by uncertainty and identification (55.3% failure), reporting and completeness (54.7% failure), biological interpretation (39.0% failure), controls and causal claims (37.2% failure), and choosing the next experiment or decision (31.0% failure). Gemini 4 Argon had the highest proportion of reviewed tasks without a recorded category failure in computation and data handling (45% success) and biological interpretation (69% success). Claude Opus 5.5 led in controls and causal claims (72% success) and choosing the next experiment or decision (75% success). GPT-6 Astra led in uncertainty and identification (56% success) which included judging the precision of estimates and recognizing when measurements could not distinguish between biological explanations. GPT-6 Astra and Meta Muse Wonder tied for the highest proportion of answers with no recorded omissions or reporting failures (51% success). Five case studies presented here show how top AI models can perform many analytical steps correctly yet misinterpret what results signify. For example, some answers described cells as fully recovered when oxygen consumption improved even though mitochondrial membrane potential and cell number remained below control levels. These model answers therefore overstated the extent of cell recovery. Other answers incorrectly called a protein a direct microRNA target, although the experiment had not tested whether the microRNA acted directly on its mRNA or affected it through another process. In a protein turnover task, some model answers misestimated protein half-lives because they didn’t account for the gradual labeling of amino acids used to make new proteins.

**Significance:** These model errors can lead researchers to pursue the wrong explanation even when individual analytical steps are completed correctly. This matters for major scientific goals such as drug discovery, where progress depends on choosing useful experiments and tools, interpreting unexpected results, troubleshooting failed approaches, and deciding which direction is most worth pursuing. Evaluating AI research assistance therefore requires checking whether its explanations, conclusions, and proposed experiments follow from the data and experimental design. These cases highlight continuing challenges in <u>scientific reasoning</u> for frontier AI models: bringing together different types of biological data, analyzing them with appropriate tools and databases, and using the results to draw appropriate conclusions and guide the next steps in research. The observed errors suggest training priorities in data analysis, biological interpretation, and experimental judgment and motivate evaluations that follow researchers and models through iterative scientific work.

## Introduction

As frontier AI models take on critical roles in scientific research, it is important to understand how well they perform the tasks researchers rely on them to do, especially as they are increasingly expected to handle biological research workflows end to end. Major goals such as drug discovery depend on many smaller dependent steps: finding relevant biological information, working with imperfect data in different formats, choosing analyses, interpreting results, and deciding which experiment to perform next. These activities connect computation with scientific decisions across basic biology research, drug development, infectious disease, environmental research, and pandemic research. Evaluating this assistance requires a clear account of strengths and weaknesses across AI model capabilities in data handling, biological interpretation, controls, uncertainty, and experimental planning.

Recent advances reflect a shared commitment across leading AI organizations to helping scientists tackle biological and medical questions. OpenAI, PerturbAI, and Scripps Research used GPT-Rosalind agents to interpret a mouse-brain CRISPR atlas and propose mechanisms for experimental investigation [1]. Google demonstrated how a researcher guided Gemini workflow in Antigravity, Google’s AI-agent workspace for connecting scientific databases and analysis tools, could help prioritize an antimicrobial peptide candidate for lab testing [2]. Anthropic used Claude to design proteins that showed experimentally confirmed binding to recombinant scorpion toxins [3]. Meta reported that Muse Spark 1.3 and Meta Muse Wonder supported medical researchers in developing hypotheses, analyzing health datasets, and preparing research manuscripts under expert direction [4]. SpaceXSI reported evaluations by LatchBio examining Grok 4.6’s capabilities in biological analysis and its safeguards in biosecurity-related tasks [5]. These efforts are accompanied by major investments in tools and infrastructure: Google DeepMind, Isomorphic Labs, and Meta announced a combined investment in Biohub’s Virtual Biology Initiative, with NVIDIA committing computing infrastructure, software, and technical expertise [6]. NVIDIA and UW Medicine’s Institute for Protein Design also released RFD4-Proteina, an open model for protein design [7]. Together, these efforts demonstrate the breadth of AI’s emerging contributions to scientific research and motivate complementary work to evaluate and improve how models analyze data, explain biological results, and help scientists choose valuable next steps.

Biology and medicine are dynamic and inherently complex fields that pose difficult challenges even for experienced scientists and physicians. Researchers work with complex living systems, limited or imperfect data, and experiments that can take weeks or months without producing a clear answer. Progress requires bringing together different types of information, choosing and using specialized tools and databases, interpreting unexpected results, and troubleshooting when an approach fails. Researchers must continually decide what a result means, which explanation best fits the observations, and which next step is worth the time and resources. Useful AI assistance therefore depends on more than biology knowledge or correct calculations: it requires applying that knowledge to the details of an experiment and helping researchers choose a productive path forward. Many valuable benchmarks have advanced the evaluation of AI across biological research activities, including literature and database use, experimental design, scientific reasoning, and molecular-data analysis [8–13]. Terminal Bench Science extends this effort across scientific disciplines through expert-authored research workflows, giving scientists a direct role in defining the capabilities they need from AI research assistants [14]. Together, these contributions provide an important foundation for understanding models’ strengths and limitations and guiding their development toward useful scientific work.

To contribute to this shared effort, we present BioResearchBench, a collection of 50 realistic research questions developed by over 100 PhD biologists drawing on their research expertise. The collection includes multi-step research tasks that require models to carry information from data processing through biological interpretation and decisions about follow-up experiments. Each question includes supporting materials, a reference answer, and a grading rubric, and reflects a problem experts would ask AI models to help address in their own research. We describe the biological areas, data types, and scientific skills represented, compare model performance, and examine how models connect their analyses to biological conclusions and proposed experiments. Five case studies show what models did well, where their answers went wrong, and how those mistakes could affect subsequent research. We consider both rubric scores and the biological meaning of individual answers, distinguishing errors that change the scientific interpretation or next experiment from other unmet requirements while recognizing useful work in responses that did not pass.

## Materials and Methods Benchmark Construction and Evaluation

### Task Development and Expert Review

PhD biologists from biotechnology, academia, and other scientific settings developed questions within their own areas of deep research expertise. Authors were asked to write realistic questions they or their colleagues would ask AI models during research. Experts were asked to develop verifiable research questions, with reference answers and supporting materials that allowed the requested outputs and conclusions to be checked. Each author prepared the question, any required input files, a reference answer, and verification materials supporting that answer. Input files were optional and varied with the biology problem. Authors also estimated how long a domain expert would need to complete each task using available tools and resources but without AI assistance. Each task underwent review by another PhD biologist and an internal PhD reviewer. Authors and reviewers exchanged feedback and revisions to improve scientific accuracy, clarity, fairness, and completeness, and to ensure that the verification materials addressed the answer (Fig. 1). Initial rubric criteria were drafted from the reference answer with AI assistance. Authors and reviewers then critically examined each criterion individually, adding, revising, or removing criteria to reflect the scientific requirements of the task. Experts were instructed to include all critical requirements they considered necessary for a correct answer. The intended grading rule required an answer to satisfy all required criteria to pass.

**Figure 1.**
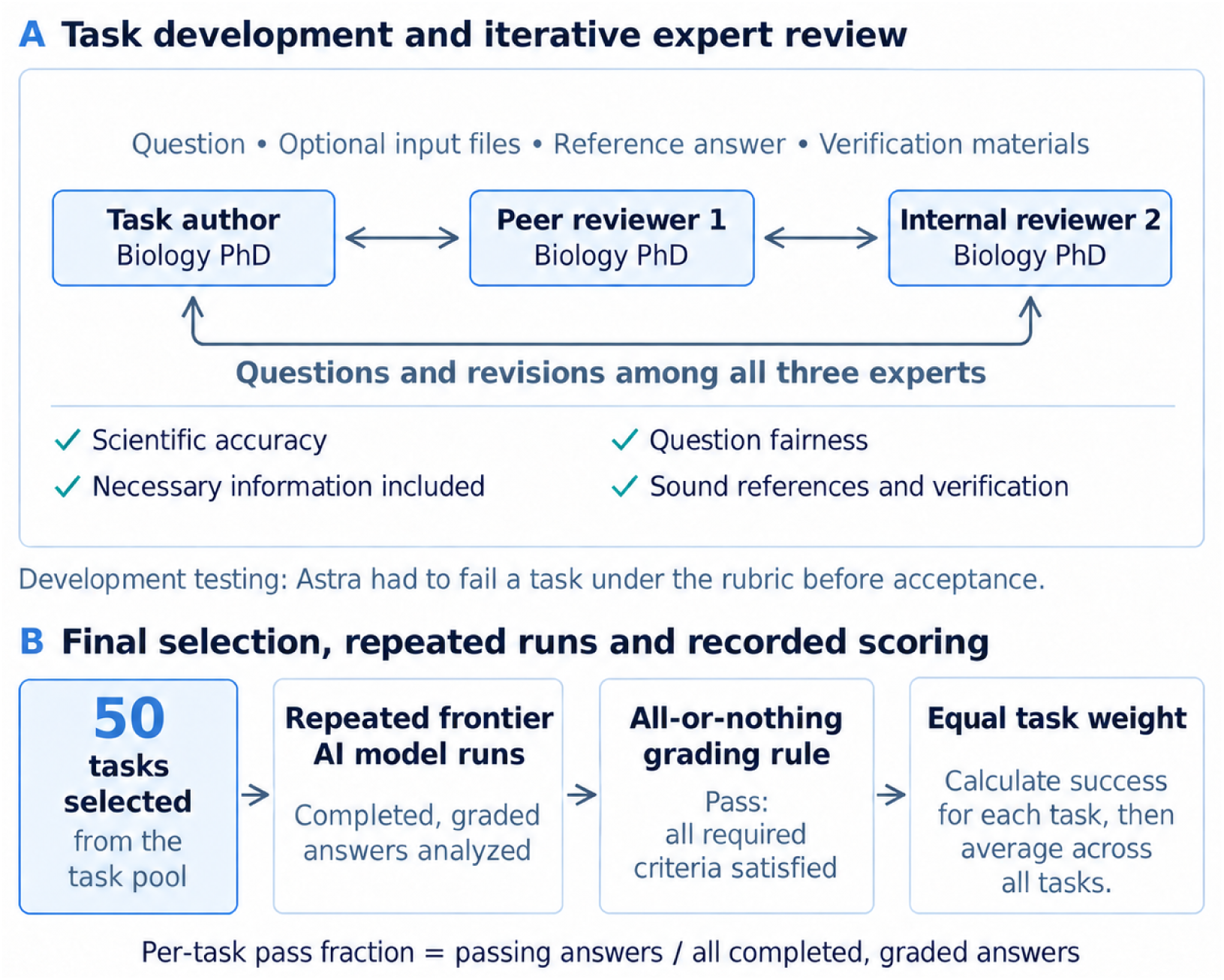
Task development and evaluation in BioResearchBench. Fifty tasks were selected for the final benchmark following development and expert review. Tasks were reviewed by another biology PhD expert and an internal biology PhD expert. Task finalization was an iterative process. Authors and reviewers exchanged questions and revisions to check that the science was accurate, the questions were fair, the necessary information was included, and the reference answers and verification procedures were sound. Task acceptance required an Astra failure under the grading rubric during development. Repeated frontier model runs produced completed, graded answers. All-or-nothing grading was intended to require every mandatory criterion to pass; the analysis retains original recorded outcomes. Each model’s pass rate is the mean of 50 per-task pass fractions computed from completed graded answers.

Task development was iterative and informed by model testing. Tasks were required to elicit a failed response from OpenAI GPT-6 Astra under the grading rubric before acceptance. This process aimed to identify realistic research problems that remained challenging and examine where model responses fell short. Some questions were subsequently revised or removed when newer models answered them correctly. The collection therefore reflects both expert review and model-informed selection for difficulty.

The final benchmark contains 50 tasks selected from this larger collection following review for scientific accuracy, clarity, and difficulty. Final selection considered biological breadth, scientific demands, and expert identified concerns about instructions, reference answers, and grading. Each task was assigned a primary biology subdomain.

### Task Package and Execution Environment

Tasks were packaged using the Harbor task format [15]. Each package separates the question, input data, execution environment, reference-solution materials, and evaluation files. The question is stored in instruction.md, configuration and metadata in task.toml, and supplied inputs within environment/data. Dockerfiles define the software environment and copy the input data into the task container. Reference materials are stored under solution/, while tests/ contains the evaluation entry point and grading rubrics. Runs were conducted offline with task inputs supplied locally and no external internet access from the solver environment. This restriction concerns the environment in which the agent analyzes data and executes commands; model inference and grading can still use services outside that environment. Run configurations specified Harbor’s Terminus 2 agent with an XML response parser and a Daytona execution environment.

### Models Time Limits and Reruns

We evaluated 5 frontier AI models: GPT-6 Astra, Claude Opus 5.5, Gemini 4 Argon, Meta Muse Wonder, and Grok 4.7. Evaluations used the Terminus 2 agent harness within Harbor allowing models to interact with the task environment, execute analysis code, and generate the requested outputs. GPT-6 Astra, Claude Opus 5.5, Meta Muse Wonder, and Grok 4.7 were evaluated using high reasoning effort. Gemini 4 Argon was evaluated at medium reasoning effort because runs at high reasoning effort encountered timeout issues. The standard execution time limit for all models was 60 minutes. These limits specify the time allowed not the time each model took to finish. Each task was run at least 4 times with each of the models. Tasks were rerun following execution or verification errors. These technical failures were recorded separately from completed responses that failed scientific grading. Success rates were calculated using only completed, graded responses excluding timeouts (Fig. S1) and any infrastructure errors.

### Grading Tasks

Model responses were assessed against expert-developed rubrics using automated LLM-based grading. The grader was Claude Opus 5. Rubric criteria were supplied together and the grader could inspect the submitted answer and generated files using filesystem tools. It returned a met-or-not-met judgment with an explanation for each criterion. Separate code-based checks verified output requirements, such as whether the required answer file existed and was nonempty; these checks did not establish biological correctness. The intended whole task passing rule required all mandatory rubric criteria and required output checks to be satisfied. Performance summaries used the recorded task outcomes and criterion-level grading results. Selected responses underwent a separate scientific review to assess the biological significance of errors and the usefulness of the analysis, as described below.

### Performance Measures

For each task and model, we divided the number of recorded passing responses by the number of eligible completed graded responses. We then averaged these fractions across the 50 tasks giving every task equal weight. Completed responses that failed the rubric remained in the denominator. Whole task success used pass/fail outcomes in the analysis dataset. Overall success rates used all eligible completed responses. The detailed scientific review and category analyses used 1 fixed, completed graded response per task and model. We selected the highest-numbered eligible attempt available at the time of selection. For the rubric-criterion failure summary, we calculated the fraction of recorded criteria failed within each selected response and averaged those fractions across the 50 tasks. Criteria were not pooled across tasks so tasks with longer rubrics did not receive greater weight. Each model received equal weight in the 5-model averages shown in figures.

### Scientific Categories and Failure Classifications

We grouped the scientific requirements in the expert-written rubrics into 6 categories: computation and data handling; biological interpretation; controls and causal claims; uncertainty and identification; next experiment or decision; and reporting and completeness. For the task overview, a task was included in a category if at least 1 mandatory criterion required that skill. The task overview counted 30 tasks with explicit reporting requirements in their rubrics. The response analysis also assessed whether answers supplied outputs requested in the task instructions, covering all 50 tasks before the reporting-specific exclusion left 49. Tasks could belong to several categories and not every task required every skill. For each selected model response, a category failure meant that the answer missed at least 1 relevant requirement even if it met other requirements in that category. When a criterion involved several skills, we classified the failure according to the component actually missed. All 5 models were compared using the same tasks within each category. Category comparisons included only tasks for which the relevant requirements and failures could be clearly classified. Two category-specific exclusions were applied consistently across all models: the glucose-metabolism task (Task ID 608) from uncertainty and the REST transcription-factor analysis task (Task ID 331) from reporting. Both tasks remained in the overall benchmark with their grades and rubric-criterion totals unchanged. The final comparisons included 49 tasks for computation, 42 for biological interpretation, 36 for controls, 43 for uncertainty, 20 for next experiments, and 49 for reporting.

### Statistics Used

We used 10,000 paired task-bootstrap resamples to describe how the summaries varied with the tasks included. Tasks, rather than individual attempts or rubric criteria, were resampled. Each sampled task retained its matched results across models with recorded outcomes, responses, and grades held fixed. Overall success and rubric-criterion failure used resamples of 50 tasks. Category-specific analyses used the corresponding eligible task set, with shared exclusions held fixed. Model summaries and equal-weight model averages were recalculated within each resample. The reported ranges were the 2.5th and 97.5th percentiles, calculated using linear interpolation and reproducible NumPy PCG64 random seeds. They describe variation in task composition within this deliberately challenging collection, not uncertainty over all possible biological research questions. They do not capture repeat-run variability on the fixed tasks, grading uncertainty, category-coding uncertainty, or selection bias. No adjustment was made for multiple comparisons and overlap between separate model ranges was not treated as a significance test. Displayed percentages were rounded to whole numbers.

### Scientific interpretation and task quality review

We reviewed the selected responses in detail alongside the task questions, available descriptions of the input measurements, reference answers, and grading records. We examined whether each analysis addressed the biology question and whether its conclusions and proposed experiments followed from the results. We distinguished claims contradicted by measurements from mechanisms that remained untested and alternative explanations that the response omitted.

## Results

### Task Identity Biology Topics and Scientific Research Skills in this Task Collection

The 50 tasks span 10 biological subdomains (Fig. 2). Genetics and physiology/metabolism are the largest groups. Cancer biology and neuroscience each account for 12% (6/50 tasks), followed by biochemistry/biophysics and immunology at 10% each (5/50 tasks), microbiology at 8% (4/50 tasks), cell/developmental biology at 4% (2/50 tasks), and medical biology and pharmacology at 2% each (1/50 tasks).

**Figure 2.**
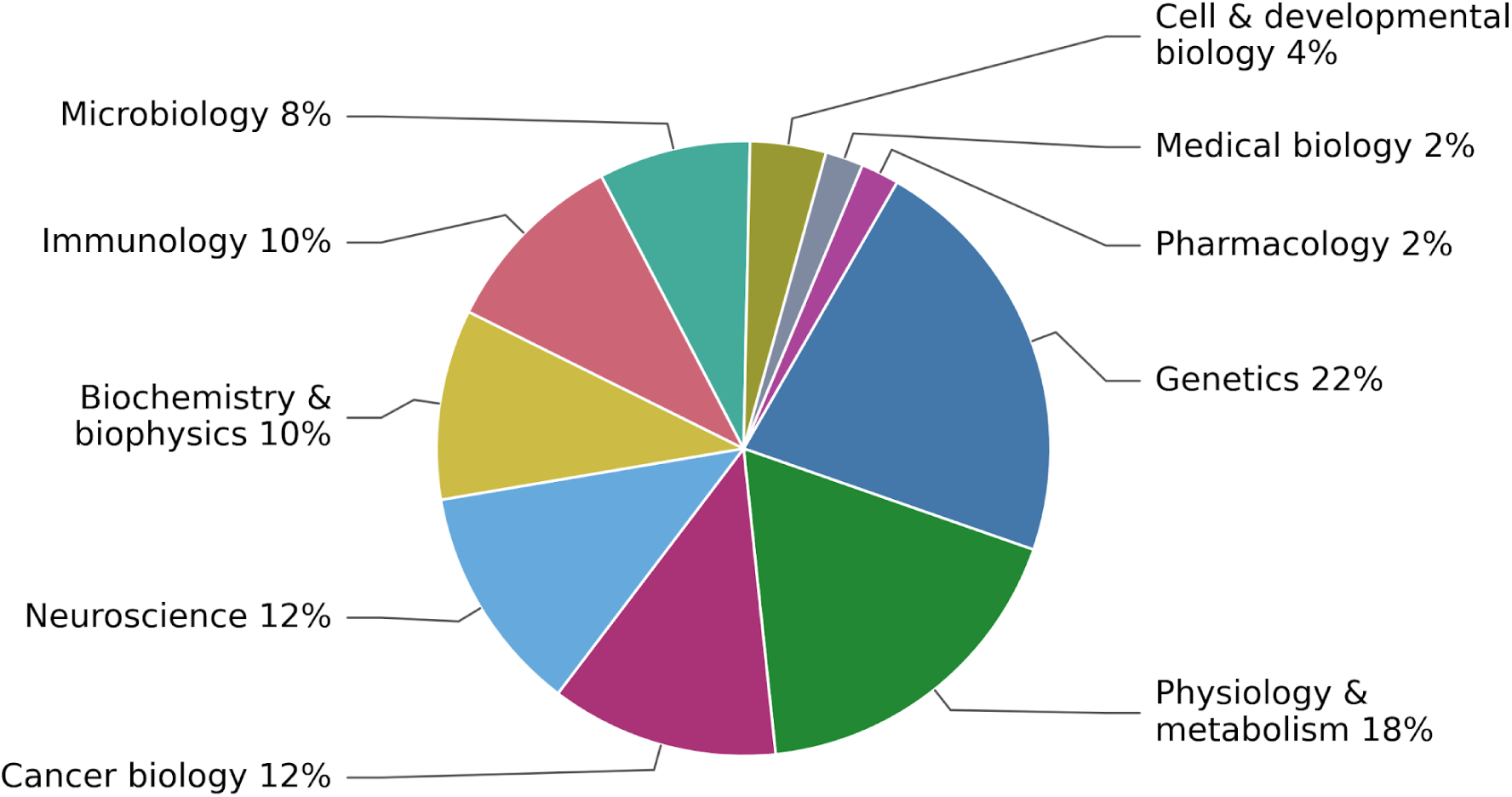
Biology Subdomains of Tasks. Areas show the percentage of the 50 tasks assigned to each primary biological domain. Each task is counted once. Genetics includes genomics, epigenetics and RNA biology; Physiology & metabolism includes metabolic regulation, circadian biology, tissue function and biomechanics; Biochemistry & biophysics includes proteomics, enzyme assays and molecular binding thermodynamics; and Cell & developmental biology includes stem-cell regulation, differentiation and skeletal phenotyping. Medical biology includes clinical tasks and Pharmacology includes receptor drug interactions and clinical adverse-event interpretation. **Significance:** The largest groups cover genetics, followed by physiology and metabolism. Cancer biology, neuroscience, immunology, and other fields are also represented.

We also examined each task’s grading rubric to identify what models were required to do scientifically (Fig. 3). Computation and data handling were required in 98% (49/50 tasks), reasoning about uncertainty or what could be determined from the available data in 88% (44/50 tasks), interpreting biological results in 84% (42/50 tasks), and evaluating controls or cause-and-effect explanations in 72% (36/50 tasks). Explicit requirements for reporting results or including particular information appeared in 60% (30/50 tasks), while 40% (20/50 tasks) required proposing a next experiment or making a decision. A single task could involve several of these requirements so the percentages overlap and do not sum to 100%. Experts reported how long it would take them to complete these tasks and generally ranged from several hours to several days. They estimated that completing their own tasks as a PhD expert without AI would require 2–20 hours for 33 tasks and 1–3 days for 17 tasks. These were the expert author estimates not measured human completion times.

**Figure 3.**
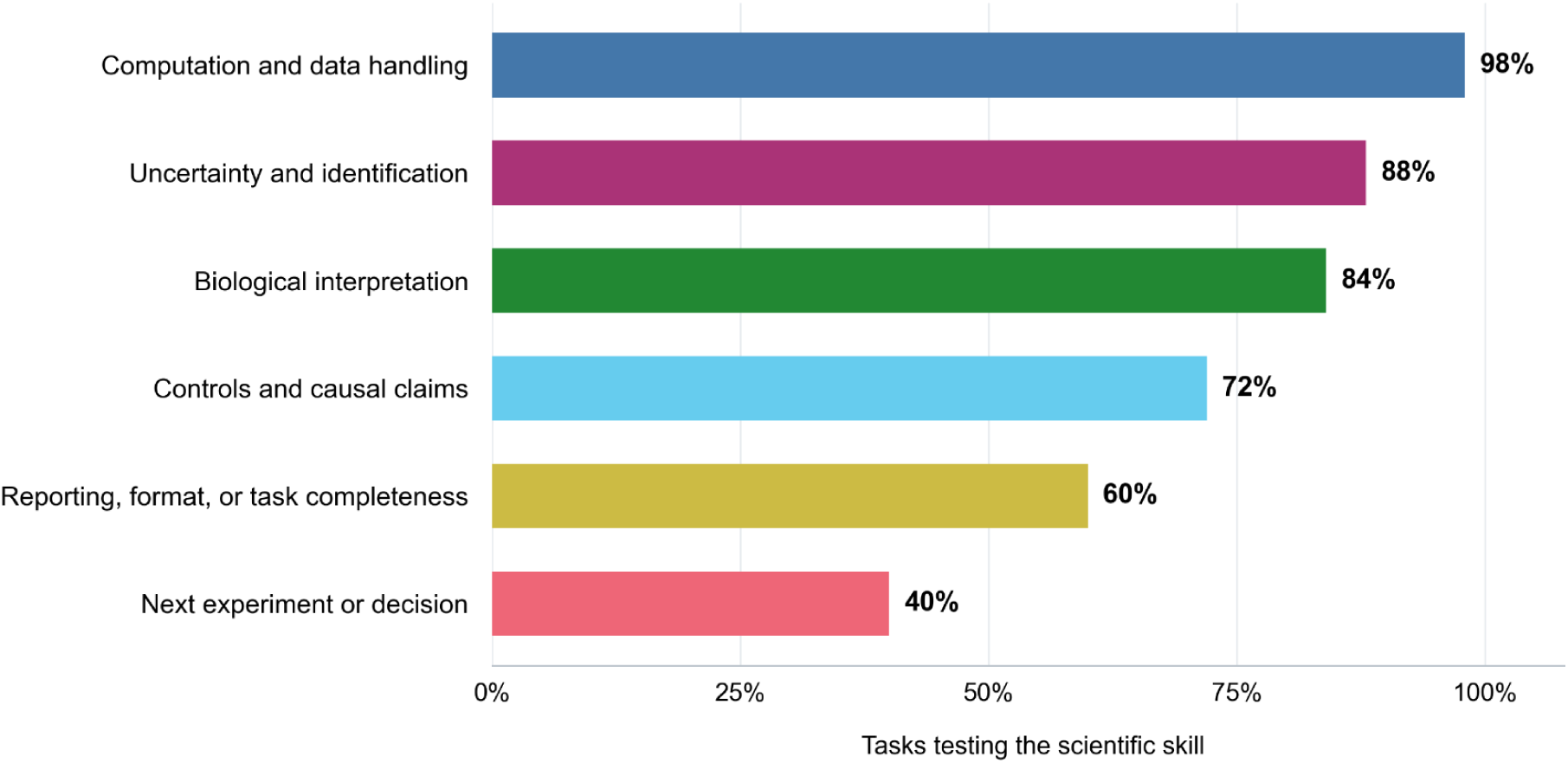
Scientific Skills Tested in the Tasks. A task contributes once to a skill when there is at least 1 mandatory rubric requirement. Among the 50 tasks, counts are 49 for computation and data handling, 44 for uncertainty and identification, 42 for biological interpretation, 36 for controls and causal claims, 30 for explicit reporting or completeness, and 20 for a next experiment or decision. Categories overlap; percentages therefore need not sum to 100%. Computation covers calculating and organizing data; biological interpretation connects measurements to the biological system; controls support causal comparisons; uncertainty includes unresolved explanations; next experiments address the remaining question; and reporting supplies the required outputs. Skill prevalence and failure eligibility use different definitions, as explained in Methods, including for uncertainty and identification and for reporting, format, or task completeness. Figure 7 uses the same taxonomy to examine model failures on these tasks. **Significance:** Almost every task requires computation, most require handling uncertainty, and some require a next experiment or decision. In practice, this can mean matching sample labels before comparing treatments, deciding what an RNA change says about protein function, or choosing a test that separates 2 explanations.

### Model Performance Across the 50 Tasks

The 50 tasks were evaluated using 5 frontier AI models: GPT-6 Astra, Claude Opus 5.5, Gemini 4 Argon, Meta Muse Wonder, and SpaceXSI Grok 4.7. Giving each task equal weight, success rates were 18% for Claude Opus 5.5, 13% for Gemini 4 Argon, 11% for Meta Muse Wonder, and 4% each for GPT-6 Astra and Grok 4.7 (Fig. 4). Success used the recorded pass/fail outcome as described in Methods. Individual task performance is shown in Figure 5. These calculations included only completed, graded responses; timeouts and infrastructure errors were reported separately and excluded from the task success calculations. These overall scores do not explain which parts of an answer were useful or why other parts failed.

**Figure 4.**
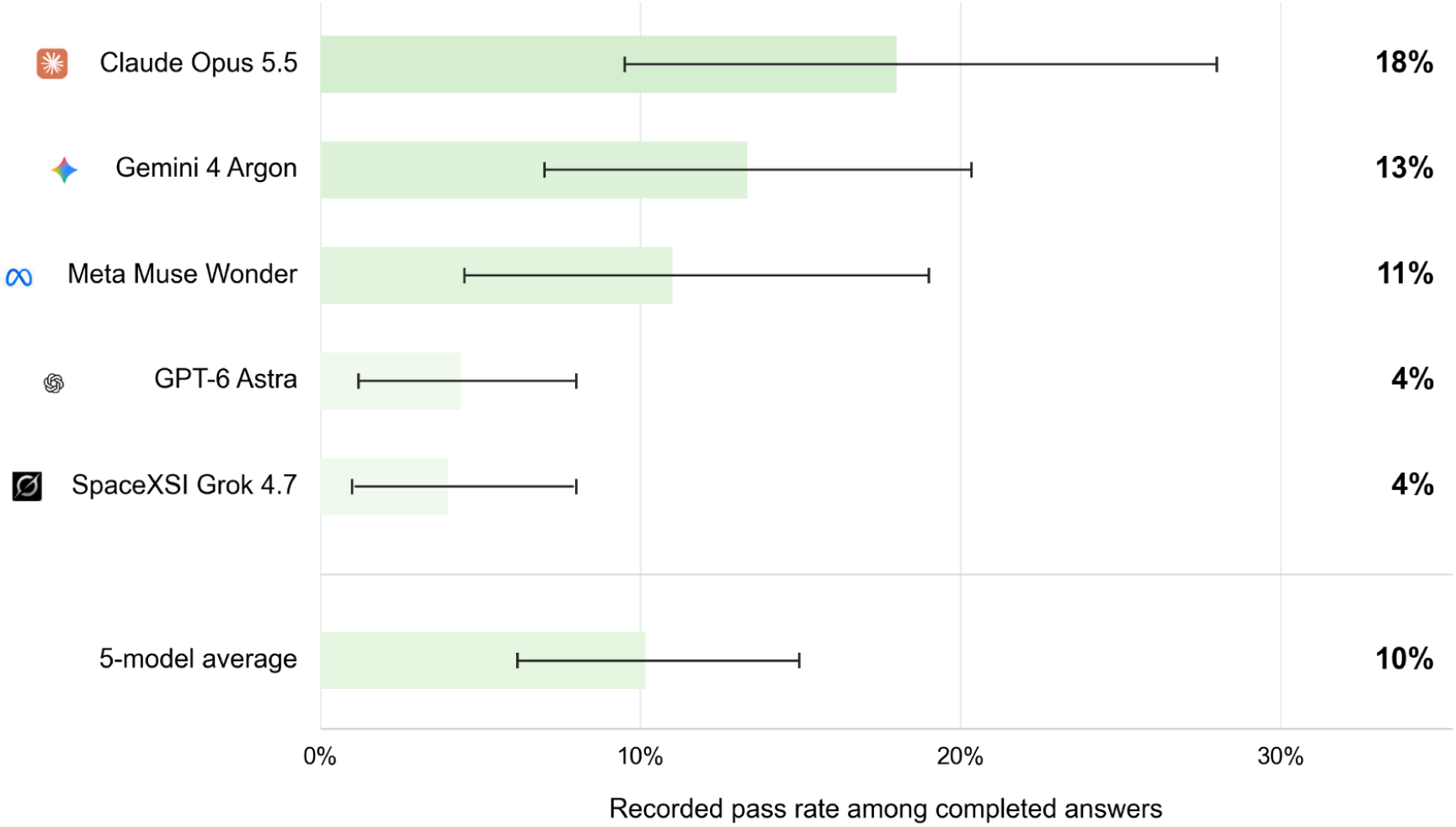
Task Success Rate. Recorded task success across 50 tasks with completed graded answers from models. Bars show task-balanced pass rates as defined in Methods. Timeouts and infrastructure errors are analyzed separately. The average bar equally averages Gemini 4 Argon, GPT-6 Astra, Claude Opus 5.5, Meta Muse Wonder, and Grok 4.7. Whiskers show the empirical 2.5th and 97.5th percentiles of 10,000 paired task resamples. These descriptive ranges are conditional on this deliberately challenging collection, whose task acceptance required an Astra failure during development. Labels show whole percentages. **Significance:** Even the highest task-balanced success rate was about 18%, so most completed answers did not receive a passing grade on this challenging set. Some failed answers correctly identified filter-induced brain signals or classified cancer-cell responses while leaving requested analyses incomplete. A low whole task score therefore motivates a closer look: which missing step changes the research conclusion and which part of the work can still be used?

**Figure 5.**
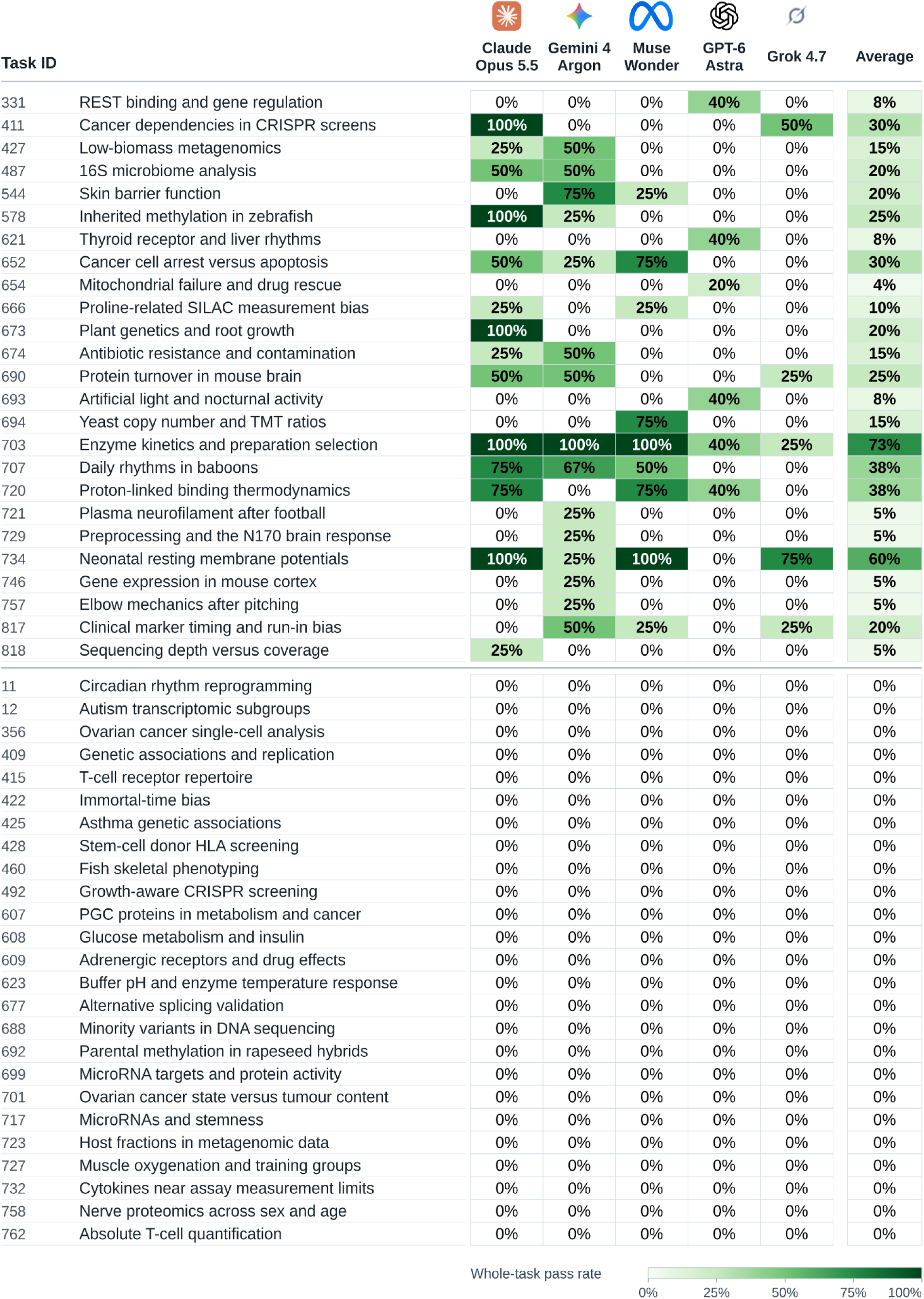
Task-level success rates. This shows passing answers divided by completed graded answers for each task and model. The final Average column is the arithmetic mean of the 5 exact model-specific pass fractions, giving each model equal weight. White indicates 0%; darker green indicates higher positive pass rates on a 0–100% scale; labels are rounded to the nearest whole percentage point. The 25 tasks with at least 1 recorded whole-task pass appear first; the 25 with no recorded whole-task passes appear below the separator. 0% indicates that none of the responses received a passing outcome for that task. **Significance:** The models did not succeed on the same problems. Twenty-five tasks had at least 1 recorded passing answer from at least 1 model, while the other 25 had none. On 11 tasks, only 1 model produced a passing answer. For example, Astra recorded passes on the mitochondria failure and drug-rescue task (Task ID 654), whereas the other 4 models did not. Opus alone recorded passes on a plant genetics and root-growth task (Task ID 673), and Meta Muse Wonder alone recorded passes on a yeast copy-number and TMT-ratio task (Task ID 694). These differences are useful starting points for reading the answers: what did one model recognize and which requirement did the others miss? They also show why an overall score does not tell a scientist how a model will handle a particular research question.

A response could fail a task while still completing much of the requested work correctly. To examine this, we also measured how many rubric requirements were missed in the selected responses. Averaged across tasks, Claude Opus 5.5 and Gemini 4 Argon each missed 18% of criteria, GPT-6 Astra missed 21%, Meta Muse Wonder missed 23%, and Grok 4.7 missed 32% (Fig. 6). Every model met most requirements on average despite low overall task success. These 2 measures describe different aspects of performance: the overall success rate records whether a response passed the task, while the criterion analysis shows how much of the required work it completed. Neither measure explains the scientific consequences of a mistake. We therefore examined which requirements models missed and how those errors affected their biology understanding and proposed next experiments.

**Figure 6.**
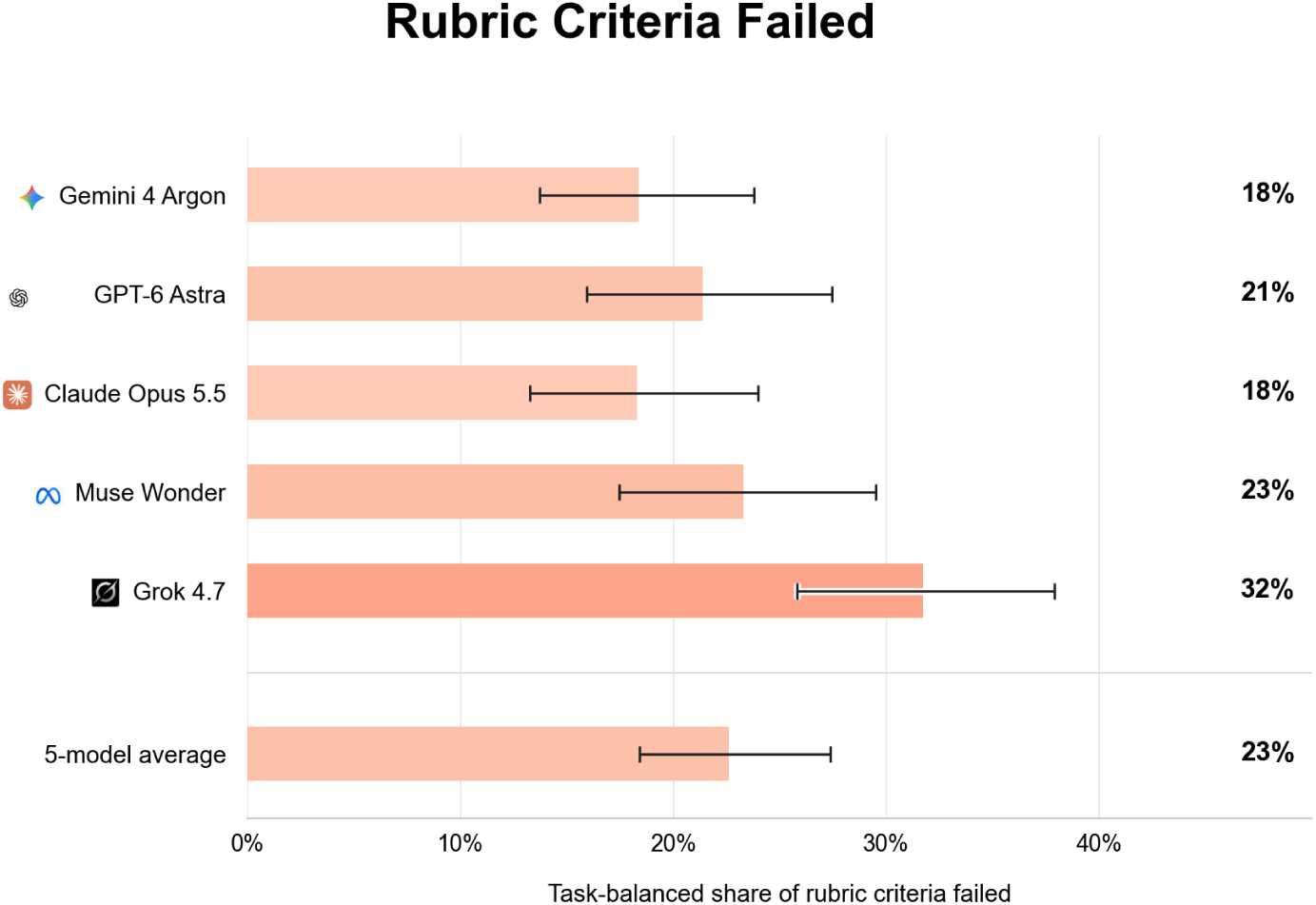
Rubric Criteria Failed. For each selected model response, the number of failed criteria is divided by the total criterion count, then averaged with equal weight per task. The bars are 18% for Gemini 4 Argon, 21% for GPT-6 Astra, 18% for Claude Opus 5.5, 23% for Meta Muse Wonder, 32% for Grok 4.7, and 23% for the 5-model average. Unlike Figure 7, which counts tasks with a category-specific failure, this figure measures the fraction of requirements missed. Labels are rounded to whole percentages. Whiskers show the central 95% of 10,000 paired task-bootstrap resamples, with the 5-model average recalculated in each (Methods). **Significance:** All 5 models fulfilled a majority of rubric criteria on average despite low whole-task success. They missed about 18–32% of criteria per task. This helps distinguish an answer that completes most of the analysis from one that leaves much of it undone.

### Models Differed in the Scientific Category Requirements They Missed

Across the 5 models, failures were most frequent in computation and data handling affecting an average of 66.9% of eligible tasks. This was followed by uncertainty and identification (55.3%), reporting and completeness (54.7%), biological interpretation (39.0%), controls and causal claims (37.2%), and next experiments or decisions (31.0%) (Fig. 7; Table 1). These percentages give each model equal weight and count tasks with at least 1 missed requirement in the category. Because the categories involve different tasks and requirements, the pattern describes this collection rather than showing that one scientific skill is inherently harder than another.

**Figure 7.**
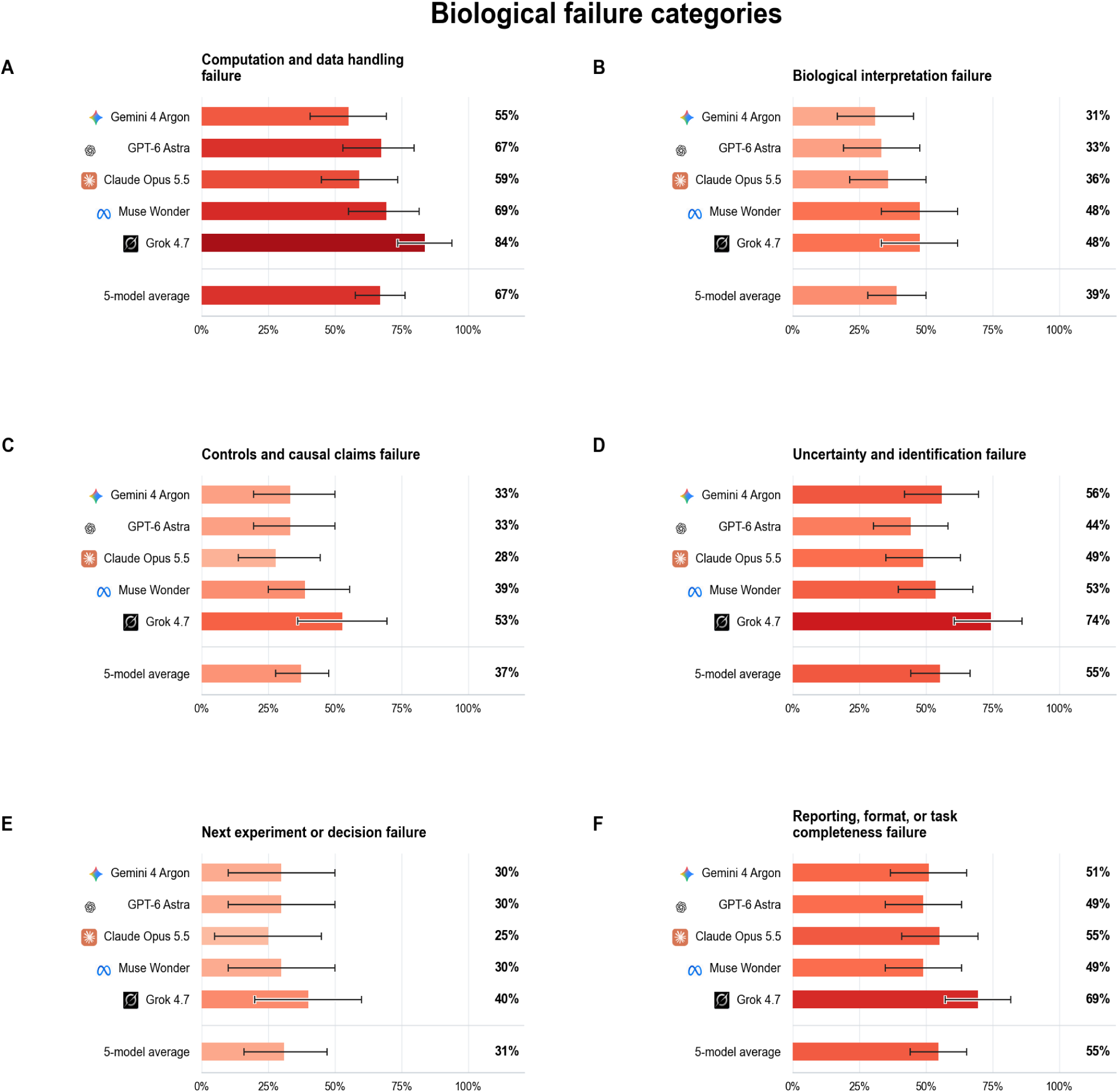
Recorded failures by scientific requirement. Each percentage is the share of tasks with at least 1 recorded failed requirement in the category using model responses to the 50 tasks. Lower values indicate fewer recorded category failures. The shared denominators are 49 for computation, 42 for biological interpretation, 36 for controls and causal claims, 43 for uncertainty and identification, 20 for next experiments or decisions, and 49 for reporting or completeness. Categories overlap; the shared exclusions are described in Methods. The average bar equally averages all 5 models. Table 1 gives exact counts. Whiskers show the central 95% of 10,000 paired task-bootstrap resamples within each category, with the shared eligibility masks held fixed (Methods). **Significance:** Models showed different strengths across the scientific categories. Gemini 4 Argon had the lowest recorded failure proportions for computation and biological interpretation, Claude Opus 5.5 for controls and next experiments and GPT-6 Astra for uncertainty. GPT-6 Astra and Meta Muse Wonder tied for the lowest failure proportion in reporting. However, these category-level results do not establish the reliability of every answer. For example, Grok 4.7 correctly accounted for gradual amino-acid labeling when estimating protein half-lives but did not identify a neighboring block of altered RNA expression in the plant-development task. Meta Muse Wonder recognized a problem with the protein-turnover interpretation but did not account for gradual labeling in its final model. Evaluating scientific assistance therefore requires checking whether a model carries its understanding of the biological problem through to its calculations, conclusions, and proposed experiments.

**Table 1.**
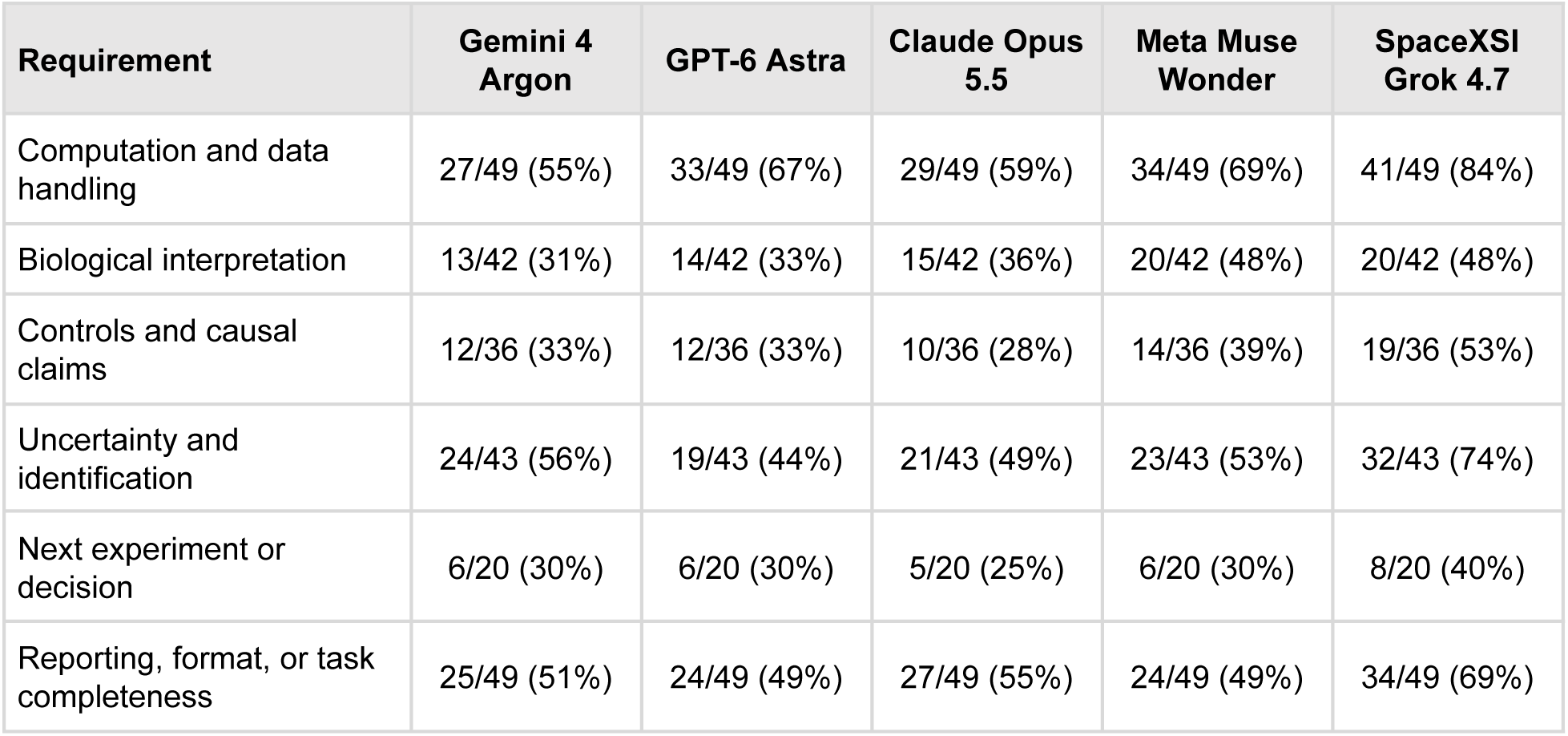
Scientific Category Failures. This table shows tasks with at least 1 attributed recorded failed component divided by category-eligible tasks, followed by the whole-number percentage. Categories overlap. Figure 7 displays the same estimates.

| **Requirement** | **Gemini 4 Argon** | **GPT-6 Astra** | **Claude Opus 5.5** | **Meta Muse Wonder** | **SpaceXSI Grok 4.7** |
|---|---|---|---|---|---|
| Computation and data handling | 27/49 (55%) | 33/49 (67%) | 29/49 (59%) | 34/49 (69%) | 41/49 (84%) |
| Biological interpretation | 13/42 (31%) | 14/42 (33%) | 15/42 (36%) | 20/42 (48%) | 20/42 (48%) |
| Controls and causal claims | 12/36 (33%) | 12/36 (33%) | 10/36 (28%) | 14/36 (39%) | 19/36 (53%) |
| Uncertainty and identification | 24/43 (56%) | 19/43 (44%) | 21/43 (49%) | 23/43 (53%) | 32/43 (74%) |
| Next experiment or decision | 6/20 (30%) | 6/20 (30%) | 5/20 (25%) | 6/20 (30%) | 8/20 (40%) |
| Reporting, format, or task completeness | 25/49 (51%) | 24/49 (49%) | 27/49 (55%) | 24/49 (49%) | 34/49 (69%) |

The model with the fewest failures differed by category. Gemini 4 Argon had the lowest failure proportions in computation and data handling at 55% (27/49 tasks), and biological interpretation, at 31% (13/42 tasks). Claude Opus 5.5 had the lowest proportions in controls and causal claims, at 28% (10/36 tasks), and next experiments or decisions, at 25% (5/20 tasks). GPT-6 Astra had the lowest proportion in uncertainty and identification, at 44% (19/43 tasks). GPT-6 Astra and Meta Muse Wonder tied in reporting and completeness, each missing requirements in 49% (24/49 tasks). Several differences between models amounted to only 1 or 2 tasks. Both GPT-6 Astra and Grok 4.7 had an overall success rate of 4%, but Grok 4.7 had higher failure proportions in all 6 categories. The largest difference was in uncertainty and identification: Grok 4.7 missed at least 1 requirement in 32 of 43 tasks (74%), compared with 19 of 43 tasks (44%) for GPT-6 Astra. Our in-depth biology review of the responses to all 50 tasks connects these differences to the biological conclusions and the experiments proposed next. We describe our results below.

The computation and data-handling category asks whether an answer obtains the requested quantity from the supplied measurements. In a task on protein turnover in mouse cortex (Task ID 690), newly synthesized proteins can contain amino-acid precursors that have not yet acquired the experimental label. Gemini 4 Argon, GPT-6 Astra, Claude Opus 5.5, and Grok 4.7 correct for this source of error when estimating protein half-lives. Meta Muse Wonder recognizes that newly made protein can remain unlabeled, but its main calculation instead treats part of the protein pool as non-exchanging. This changes the biological explanation for the unlabeled signal and produces different half-life estimates and comparisons between mouse groups. The correction matters because otherwise newly made protein can be counted as old protein making replacement appear slower than it is. Biological interpretation asks what a result means in the biological system being studied. In a task on microRNA regulation (Task ID 699), the protein assay measures activity-associated enrichment rather than protein abundance directly. A reduced signal together with a matching microRNA-binding sequence identifies a candidate for further testing. Gemini 4 Argon instead uses labels that imply direct regulation has already been validated, although that experiment has not been performed. The controls and causal-claims category asks whether the experimental comparisons separate the proposed cause from other explanations. In a cancer-cell dependency screen (Task ID 411), several CRISPR guides targeting the same amplified genomic region can all reduce cell growth because cutting many DNA copies is toxic. Agreement among guides therefore leaves the gene’s contribution to growth unresolved. Reducing gene function without cutting DNA would test that explanation more directly. The distinction between different types of control is illustrated by a task on sphere formation by cultured mammary epithelial cells, a breast-cell model, after reducing the regulatory RNA miR-424 (Task ID 717). The assay measures the formation of cell clusters called spheres. The task asks whether reducing other regulatory RNAs would provide useful controls. Grok 4.7 rejects reducing miR-145 as a comparator because it would not be neutral even though it correctly notes the expected increase in stem-like activity. A positive control is deliberately expected to change the readout: reducing miR-145 could test whether the assay detects increased sphere-forming activity. Whether miR-424 specifically caused a response is a separate question. The error is treating every useful control as a comparison that should leave the system unchanged. The uncertainty and identification category asks both how precise an estimate is and whether the available observations determine the quantity or explanation being requested. In a cortical protein-turnover task (Task ID 690), some time courses do not fit the model based on the measured cortical precursor. Several explanations could account for that mismatch so assigning another tissue as the protein’s source requires additional data. GPT-6 Astra preserves this distinction more consistently than Claude Opus 5.5. The next experiment or decision category asks whether the proposed action would address the remaining biological question. In an Arabidopsis root development task (Task ID 673), a linked genetic change could explain part of the expression pattern attributed to ORE15. An independent mutant or a line with restored ORE15 function would help test that explanation before a candidate binding list is interpreted as evidence of direct regulation. The reporting and completeness category asks whether an answer supplies the requested analyses, values, comparisons, and other outputs. In an enzyme-kinetics task, GPT-6 Astra appropriately keeps its conclusion conditional but omits a numerical sensitivity calculation identified in the recorded grading explanation. The requested calculation is needed to assess how sensitive the conclusion is to the assumptions. The final answer should state consistently which results were obtained and which questions remain open as this is a vital part of biology research.

### Models Reached Correct Biological Conclusions Despite Errors in the Analysis

The overall scores do not show differences between an answer that misses the central biological point and one that reaches a useful conclusion but leaves other work incomplete or incorrect. Three additional tasks illustrate this distinction before the 5 detailed case studies we present later. In a brain recordings task, all 5 models recognized a processing artifact but some omitted requested analyses. In a cancer cells task, the models agreed on broad drug response patterns while some supporting calculations remained inaccurate. In an activated T cells task, useful RNA–protein comparisons were accompanied by missed quantitative corrections and in some answers, overly specific explanations. These observations come from comparing selected responses with expert ground truth answers.

### Models Recognized a Brain Recording Artifact but Left Parts of the Analysis Incomplete

A task on human brain responses to visual stimuli (Task ID 729) asks whether an early electrical signal reported at about 88 milliseconds reflects the brain’s response to faces or an artifact from processing the recording. Electroencephalography (EEG) records electrical signals through electrodes. Filtering removes parts of the signal but can also change the shape of a brief response. The supplied data include averaged responses to faces and houses from 24 participants at 4 electrodes, recordings processed with 0.01 Hz and 1.0 Hz high-pass filters, a waveform figure, and a report documenting a 17 ms delay before the picture appeared. The higher filter setting removes a broader range of slow signal components. Models must compare the processing choices, account for the display delay and report response sizes, timing, and statistical tests.

All 5 models’ answers identify the early positive signal as an effect introduced by the higher filter setting and apply the display delay correction. Gemini 4 Argon, GPT-6 Astra, and Claude Opus 5.5 also retain the N170, a later electrical response sensitive to faces. This is an important interpretation: changing how the same recordings are processed changes the early feature so that feature should not be reported as a new neural response to faces. The answers also differ in completeness. Meta Muse Wonder and Grok 4.7 omit requested details about a later response and the effect of filtering on comparisons across electrodes. Recognizing the early artifact therefore represents a shared success within answers that did not complete every requested analysis. The correct central interpretation and the remaining omissions should be considered separately.

### Models Distinguished Drug Responses Despite Errors in the Supporting Calculations

A task on drug-induced changes in colorectal cancer cells (Task ID 652) asks how 4 compounds reduce cell number in 2 cell lines, HCT116 and HT-29, and whether 1 candidate affects either line more strongly. A lower cell count could result from slower division, greater cell death, or both. The task therefore requires combining cell counts with measurements that reveal what the cells are doing. Eight CSV files describe 3 paired experiments, including counts at the start and after 72 hours, DNA content and synthesis, cell-death-associated caspase activity, live-imaging records, and information about plates, transfers, and staining. Models must resolve labeling and quality-control problems, calculate growth relative to the starting counts while accounting for each cell line’s untreated growth, and interpret the compounds’ effects.

All models identify a compound associated with G1 arrest in which cells stop progressing before copying their DNA. They distinguish it from compounds associated with apoptosis, a regulated cell-death process; G2/M arrest, a block after DNA copying or during division; and weak activity. All conclude that the data provide insufficient evidence that the nominated candidate preferentially affects either cell line. These conclusions agree with the expert ground truth answer. The models use the additional measurements to separate reduced cell division from increased cell death rather than interpreting every decrease in cell number as the same biological response. Some supporting analyses still require correction. Meta Muse Wonder notes the limited power of the comparison between cell lines. Grok 4.7 incompletely removes plate-edge fields and reports some growth response estimates outside the reference tolerances. The broad biology classifications remain consistent with the reference, but the numerical estimates and quality-control decisions are wrong. This example separates recognizing the type of drug response from accurately measuring its magnitude.

### Models Detected RNA Protein Differences but Missed Corrections Needed to Quantify Them

A task on helper T-cell activation (Task ID 762) asks how RNA and protein quantities change when immune cells are stimulated. It uses synthetic measurements representing initially unactivated, or naive, CD4 T cells from 6 donors before and after 48 hours of anti-CD3/CD28 stimulation. Nine files provide RNA counts, added reference molecules called spike-ins, cell counts and calibration records, peptide measurements used to estimate proteins, sample assignments, and DNA-content information. The analysis must estimate RNA and relative protein quantities per cell. A larger total signal could reflect more cells rather than more material in each cell, so calibration and cell counts are essential.

Models must classify 29 candidate genes, revisit 8 biological conclusions, and propose a confirming experiment. Gemini 4 Argon, GPT-6 Astra, and Claude Opus 5.5 identify important differences between RNA and protein changes. For SLC2A1 and ACACA, protein quantities increase while RNA remains in the task’s unchanged class. For IL2 and IFNG, they distinguish insufficient protein evidence from an observed decrease. These qualitative conclusions agree with the reference. However, all 3 answers omit the correction for RNA reads originating from other samples, leaving errors in the numerical estimates even when particular biological classifications are correct.

Meta Muse Wonder recovers the overall per-cell expansion and identifies several sample and calibration problems, but also misses the correction for RNA reads assigned across libraries and labels a relative protein scale as protein mass. Grok 4.7 recovers the large increase in RNA per cell but likewise labels its protein scale as mass and misses specific calibration, digestion, and cross-library read-assignment problems.

The proposed follow-up for SLC2A1 illustrates a separate challenge: deciding which experiment would explain an increase in protein abundance. SLC2A1 encodes GLUT1, a transporter that helps activated T cells take up glucose to support growth and immune function [16]. More GLUT1 protein without a corresponding RNA increase could arise from faster protein production, slower breakdown, or both.

Meta Muse Wonder proposes polysome profiling, which examines how RNA is associated with ribosomes—the structures that make proteins—and says that finding more ribosomes on SLC2A1 RNA would confirm increased translation. More ribosomes can also accumulate because they move along the RNA more slowly, so their increased association with RNA does not by itself show that more protein is being produced [17]. A pulse–chase experiment, which labels newly made protein and follows its disappearance, could help distinguish faster production from slower breakdown. This would help identify how activated immune cells raise GLUT1 levels, rather than directing further work prematurely toward protein production alone.

### 5 Biology Task Case Studies

We next examine 5 research question tasks in greater biological detail, tracing how the models analyzed the supplied data, where their interpretations differed, and how those differences affected the proposed next experiment. Table 2 summarizes the main findings across these cases.

**Table 2.** Biology consequences of different interpretations across 5 tasks. The table compares biological conclusions and proposed next experiments from the reviewed model responses. Detailed Results paragraphs explain the input measurements, successful analyses, and remaining problems in each case.

| **Task ID** | **Differences in model responses** | **Research Implications** |
|---|---|---|
| Cancer dependency<br>411 | All 5 account for high-copy cutting. GPT-6 Astra, Gemini 4 Argon, Meta Muse Wonder, and Grok 4.7 withhold affected calls; Claude Opus 5.5 adjusts the shared effect. | Reduced growth may result from DNA-cutting damage rather than loss of a required gene. Test the gene using a method that does not cut DNA, and check whether restoring its function rescues growth. |
| Cortical turnover<br>690 | 4 models apply the precursor correction. Meta Muse Wonder instead headlines a non-exchanging-pool model. Corrected calculations can still overstate tissue origin or treatment effects. | New proteins can contain unlabeled lysine, so the remaining light signal includes old and newly made protein. Measure precursor labeling and add earlier sampling to assess rapid replacement. |
| Respiratory rescue<br>654 | GPT-6 Astra, Claude Opus 5.5, and Grok 4.7 preserve incomplete recovery. Meta Muse Wonder’s final recovery statement conflicts with its table; several answers overstate the molecular route. | Respiration improves more than cell number or membrane potential. Report these outcomes separately and directly test the proposed substrate-transport or metabolic explanation. |
| MicroRNA targeting<br>699 | 4 models identify PREPL and ACOT1; Grok 4.7 omits PREPL from its written shortlist. Strong direct-target claims and incomplete screening checks weaken some shortlists. | The screens identify candidates for further testing. Alter the predicted microRNA-recognition site in a reporter and measure protein abundance and enzyme activity separately to test the regulatory mechanism. |
| Root development<br>673 | 4 models exclude the linked expression block; Grok 4.7 includes affected genes. Meta Muse Wonder excludes the block but overstates some technical explanations. GPT-6 Astra treats its explanation as something to test, but its proposed follow-up experiment leaves out the alternative that the plant’s carbon status contributes to the shorter roots. | The RNA changes in neighboring genes may reflect a linked DNA-copy change rather than regulation by ORE15. Check the genomic DNA and compare an independent mutant or a line with restored ORE15 function. |

### Cancer Cell Dependency Screening Task Separating Gene Dependence from DNA Cutting Damage Task ID 411

This task asks which genes cancer cells need to keep growing (Figure S2). The experiment uses a pooled CRISPR screen in which different cells receive guides that direct a DNA-cutting system to different genes. If disrupting a gene slows growth or kills cells, cells carrying guides to that gene become less common over the 21-day experiment. The models must use these changes to identify candidate growth dependencies: genes on which the cancer cells may depend. The supplied data include guide counts from the starting plasmid pool and 4 endpoint cultures, guide-cutting locations, reference sets of essential and nonessential genes, and the number of DNA copies in different genomic regions. Copy number matters because cancer cells can carry many copies of a chromosome segment. Cutting many copies can itself harm the cell, even when the targeted gene is not required for growth. A decrease in cells carrying a particular guide can reflect either loss of an important gene or damage from cutting DNA. Several guides aimed at the same amplified region can share this problem. All 5 model answers exclude 1 endpoint culture whose results disagree with the others and recognize the problem posed by high-copy regions. Agreement among guides usually strengthens a screen result, but the 4 guides targeting ERBB2 can all cause damage by cutting its amplified region. Their agreement therefore leaves 2 explanations for reduced growth: the cells depend on ERBB2 or they are harmed by the repeated DNA cuts.

The models differ in how they handle this uncertainty. GPT-6 Astra withholds dependency calls for amplified genes because the screen cannot reliably separate gene loss from cutting damage. Gemini 4 Argon also excludes high-copy regions, using a different threshold. Claude Opus 5.5 instead estimates the cutting effect shared within each region and adjusts the results, removing affected dependency calls. It also notes that removing a call leaves the gene’s contribution to growth unresolved. The distinction is between withholding a conclusion where the assay is difficult to interpret and attempting to recover a gene-specific signal by correcting for the regional cutting effect. Meta Muse Wonder also removes the discordant culture and excludes high-copy regions treating ERBB2 and CDK4 as untestable rather than essential. It requires at least 3 retained guides for a gene-level call leaving more genes unanswered than the reference analysis. Grok 4.7 uses a 2-guide minimum while accounting for high-copy cutting effects. The 2-guide rule retains genes that have only 2 usable guides, whereas the 3-guide rule excludes them. This difference changes the coverage of the gene list; the comparison does not show that either threshold produces more accurate dependency calls.

All models recognize how the screening method can create an apparent gene dependency. Their differences concern which genes they leave unanswered, whether they attempt a correction, and how they choose eligibility rules. A withheld or removed dependency call leaves open whether the gene is biologically important; it indicates that the screen has not provided a reliable answer under that analysis. The next experiment must address the source of uncertainty. Reducing the candidate gene’s function without cutting DNA would help test whether the growth effect reflects the gene itself. In that experiment, restoring gene function would provide a further check that loss of the gene caused the response. Testing neighboring amplified genes would also help assess whether the effect is specific to the candidate. Separately, verifying the identity of the discordant culture would address the replicate discrepancy. These follow-ups answer different questions: one investigates whether the cultures are comparable while the other separates a gene’s biological role from damage caused by the screening method.

### Mouse Brain Protein Turnover Task Accounting for Labeling Before Interpreting Protein Replacement Task ID 690

This task asks how quickly proteins are replaced in the mouse cortex, the brain’s outer region, and how this differs between ordinary mice and the genetically altered 5xFAD model (Figure S3). Proteins are continually made and broken down. The experiment follows their replacement using labeled lysine, an amino-acid building block that can be detected after incorporation into newly made protein. The models must estimate protein half-lives, the time needed for half of a protein pool to be replaced, and use the results to compare the 2 groups. The supplied data report the fraction of each protein that remains unlabeled, together with labeling of the free lysine available for protein synthesis, at days 4, 8, 16, and 32. Different mice provide tissue at each time point. The important complication is that newly made protein can remain unlabeled if the lysine used to make it has not yet acquired the label. Counting all unlabeled protein as old protein would make replacement appear slower than it is. The calculation must therefore account for labeling of the precursor pool. The half-life estimates also depend on approximately stable protein abundance and on the measured free lysine representing the lysine used for synthesis.

GPT-6 Astra, Claude Opus 5.5, Gemini 4 Argon, and Grok 4.7 account for precursor labeling. This is a meaningful success because the correction separates delayed labeling of the building blocks from slow replacement of the protein itself. Meta Muse Wonder recognizes the labeling issue in its explanation but uses a model with a non-exchanging protein fraction for its main estimates. Its selected response misses criterion 3: “The response estimates half-lives with a precursor-product kinetic model that uses the free lysine light fraction as the precursor (for example dL/dt = k (P(t) - L)); any equivalent precursor-corrected model is accepted.” This is more than a different numerical method: it changes the biological explanation for why protein remains unlabeled altering the estimated half-lives and the comparison between mouse groups. Even after the labeling correction, some proteins remain difficult to interpret. Claude Opus 5.5 distinguishes proteins with measurable half-lives, proteins replaced too quickly for the sampling times to provide more than an upper limit when the model fits, and proteins whose time courses do not fit the measured precursor pool. GPT-6 Astra is more cautious than the reference about some limits in ordinary mice. That disagreement concerns whether the labeling assumptions fit the observations not failure to perform the correction.

This is especially important when a protein’s labeling pattern differs from the measured cortical precursor. Claude Opus 5.5 interprets such patterns as evidence that proteins were made outside the cortex. However, other explanations remain possible, including a different precursor pool within the tissue, a mixture of protein pools, or measurement error. Gemini 4 Argon reports formal limits for some incompatible time courses and a ratio that it also acknowledges is not a valid comparison of brain tissue itself. GPT-6 Astra more consistently treats a poor fit as a reason to investigate the assumptions before assigning a tissue of origin. That restraint is useful because it avoids turning a mismatch in a mathematical model into a claim about where a protein was produced.

Planning a treatment experiment introduces a separate problem. Meta Muse Wonder uses a genotype difference in Apoe as an effect size for a drug study. Grok 4.7 correctly applies the precursor model but also uses the genotype difference in day-8 labeling to plan a drug study. Both selected responses miss criterion 18: “The response states that the WT versus 5xFAD difference is not an estimate of the antibody’s treatment effect, so the power calculation needs an assumed treatment effect on a measurable endpoint (for example the Apoe light fraction at a fixed day) together with the variance between mice.” A difference between ordinary and genetically altered mice describes the comparison already measured. It does not tell the researcher how strongly the proposed antibody will change the outcome. Using that difference as the expected treatment effect can therefore produce a sample-size calculation based on the wrong biological comparison.

The models also differ in how specifically they explain protein location. Meta Muse Wonder assigns a liver source or plaque-associated pool to some signals while Grok 4.7 interprets a fitted plateau as protein retained in plaques. These explanations require measurements of origin or location. The labeling curves leave those questions open. Thus, using the correct turnover calculation is an important first step, but the interpretation still depends on what was actually measured. The remaining questions call for different follow-up experiments. Earlier sampling would help estimate rapid turnover. Comparing precursor labeling in plasma and cortex, together with documenting how blood was removed from the tissue, would help investigate an outside contribution to the signal. A treatment pilot would provide information about the expected drug effect and variation between mice on a measurable endpoint. Finally, replacement of complement proteins, which participate in immune responses, is different from their activation; an activation assay would be needed to address that question. The main finding is that correcting the labeling model improves the half-life estimate, while judgments about tissue origin, treatment effects, and immune activity require additional biological data.

### Mitochondrial Recovery Task Separating Improved Respiration from Complete Cellular Recovery Task ID 654

This task asks why treatment with NRX13 reduces mitochondrial respiration and why adding methyl pyruvate restores some of it (Figure S4). Mitochondria use nutrients and oxygen to help produce ATP which supplies energy for cell work. Reduced oxygen consumption could reflect fewer cells, less mitochondrial material, impaired mitochondrial function, or insufficient usable fuel. The models must bring together several measurements to determine which explanations fit the observations and what remains uncertain. Seven CSV files combine oxygen-consumption measurements from a Seahorse instrument with sample-tracking records, microscopy, ATP and protein measurements, and proteomic data. Samples from 6 donors are compared across matched vehicle-control, NRX13, NRX13 plus methyl pyruvate, and rotenone conditions. The vehicle control receives the treatment carrier without NRX13. Comparing treatments within donors preserves the experimental pairing; treating every well or image as a separate donor would exaggerate the amount of independent information. The way each measurement is normalized changes the biological question it answers. Oxygen consumption per well describes the culture’s total activity, whereas consumption per cell describes the average remaining cell. Normalization to mitochondrial material asks how respiration relates to the amount of that material. A well can contain fewer cells but retain a similar total amount of mitochondrial material if each remaining cell contains more of it. Improved respiration per cell therefore needs to be considered alongside cell abundance and other measures of function when judging recovery.

Gemini 4 Argon, GPT-6 Astra, and Claude Opus 5.5 correct a sample or plate-layout swap and an image-area unit error. They retain within-donor comparisons and recognize that fewer cells coexist with preserved total mitochondrial material. Meta Muse Wonder also corrects sample identities and image units and compares treatments within donors. These are useful analytical successes: the models avoid explaining reduced oxygen consumption solely as a loss of mitochondrial material and identify a pattern consistent with limited fuel entry or utilization. The answers differ more sharply when describing how fully the cells recover. GPT-6 Astra keeps recovery incomplete and notes that pyruvate transport and pyruvate dehydrogenase (PDH) activity were not directly measured. PDH is an enzyme involved in converting pyruvate into a form that can enter mitochondrial energy metabolism. Claude Opus 5.5 also reports that cell abundance and mitochondrial membrane potential, the electrical difference across the mitochondrial membrane, remain below control levels. Grok 4.7 likewise retains the residual membrane-potential deficit. These distinctions are biologically important because improved oxygen consumption can occur while other measured properties remain impaired. Gemini 4 Argon describes near-complete respiratory recovery but broadens this to complete functional rescue in its final table. Meta Muse Wonder’s final recovery statement conflicts with its own numerical results: it states that all functional rescue intervals include 0, although its reported membrane-potential deficit is −0.053, with an interval from −0.076 to −0.029. That interval remains below 0 and indicates a residual deficit in this readout. Both selected responses miss the rubric’s recovery requirement (criterion 14: “NRX13_MePyr must be classified as a strong but incomplete functional rescue.”). The error is not overlooking the improvement; it is extending that improvement to cellular functions that had not recovered to control levels.

Identifying the affected molecular mechanism presents a separate challenge. Claude Opus 5.5 describes a specific defect in pyruvate entry and a bypass route more strongly than the measurements justify. Gemini 4 Argon and Meta Muse Wonder also make overly specific claims about transport or pyruvate metabolism. Their selected responses miss criterion 15: “The response must keep the causal claim within the data. The dataset supports a substrate entry or utilization phenotype and functional bypass or compensation, but it does not directly prove mitochondrial pyruvate transport, PDH activity, a specific MePyr entry route, or an exact molecular causal step.” Adding methyl pyruvate provides information about how the cells respond to an alternative fuel source, but several affected steps could produce that response. Direct transport or enzyme measurements are needed to identify the particular step affected by NRX13. A shared image-processing error affects the supporting analysis. All 5 selected responses exclude an entire image field after identifying a problem in the computer’s detection of cell nuclei. The defect concerns the nuclear measurement, while other channels in the same field remain usable. All 5 miss criterion 3: “The response must treat E08 field 4 as a nuclear segmentation failure only. Its nuclei based cell estimate is excluded, while mitochondrial area, TMRM, and TOMM20 from that field are retained. Excluding the whole field fails.” TMRM provides a readout related to mitochondrial membrane potential, while TOMM20 is a mitochondrial protein marker. Removing all channels discards measurements that were not shown to be defective. GPT-6 Astra and Gemini 4 Argon also examine exclusion of only the nuclear measurements and report small changes, providing a useful sensitivity check despite the recorded failure. This case separates 3 judgments that a single recovery label can obscure: whether respiration improves, whether the other measured cellular properties recover, and whether the treatment response identifies the drug’s molecular action. GPT-6 Astra, Claude Opus 5.5, and Grok 4.7 preserve incomplete recovery, whereas Gemini 4 Argon and Meta Muse Wonder overstate it. Several answers also move from a fuel-related response to a specific molecular explanation without the corresponding measurements. Follow-up experiments should therefore report recovery separately for respiration, membrane potential, and cell abundance, and directly test transport or enzyme activity before assigning a molecular mechanism.

### MicroRNA Regulation Task Identifying Candidate Targets Without Mistaking Them for Validated Targets Task ID 699

This task asks which proteins may be directly regulated by miR-185 in a lab grown cell line called Huh7.5 (Figure S5). MicroRNAs are short RNAs that can bind matching sequences in messenger RNAs and reduce the protein made from them. The task combines 2 observations: a protein-associated assay signal that decreases after adding miR-185, and a matching sequence in the candidate’s messenger RNA. The supplied files contain protein signals, sequences from messenger RNA 3′ untranslated regions, and the miR-185 sequence. The untranslated region lies outside the protein-coding sequence and can contain regulatory binding sites. Models must compare the assay signals, find sequence matches, and produce a candidate list with binding-site locations.

Understanding what the protein assay measures is essential. The FP-biotin probe captures activity-associated signal from a class of enzymes called serine hydrolases. A lower signal can reflect less protein, lower enzyme activity, or less efficient probe capture. It is therefore not a direct measurement of total protein abundance. The experiment compares control microRNA with miR-185, uses isotope labels to distinguish samples, and includes an untransfected sample without the capture probe as a background check. Nine runs belong to 3 replicate blocks. Comparisons must preserve that grouping so repeated measurements are not treated as independent biological replicates.

Gemini 4 Argon, GPT-6 Astra, Claude Opus 5.5, and Meta Muse Wonder identify PREPL and ACOT1 as candidates supported by both the protein-signal and sequence analyses. PREPL has an 8-nucleotide matching site and ACOT1 a 7-nucleotide site of the 7mer-m8 class. These sites match the microRNA seed, the short sequence that helps determine which messenger RNAs it recognizes. Finding both a matching site and a decreased assay signal gives a biological reason to prioritize a candidate, but direct action through that site still requires testing. GPT-6 Astra retains matched comparisons within each replicate block and applies Benjamini–Hochberg correction for testing many proteins. No protein meets its chosen significance threshold after correction. It therefore presents the candidates as exploratory and its combined score as a way to rank them, rather than a probability of direct regulation. This is useful restraint: a candidate can merit further testing even when the analysis does not support a confident individual discovery. However, GPT-6 Astra declines to impose a cutoff using the sparse, untransfected background sample, leaving the rubric’s probe-dependence requirement unmet. Claude Opus 5.5 checks whether the signal depends on the capture probe, summarizes matched ratios within blocks, and requires a decrease in every block. It also describes PREPL and ACOT1 as candidates. These checks address different questions: probe dependence helps assess whether the captured signal is specific to the assay, while consistency across blocks assesses whether the observed change is reproducible. Agreement on a shortlist therefore conceals differences in how thoroughly the models evaluate the measurements supporting it.

Gemini 4 Argon adds sequence-context and RNA-isoform features, but describes candidates as validated direct targets without an experiment testing action through the proposed binding site. Its answer also leaves the probe-dependence filter unapplied. Gemini 4 Argon and GPT-6 Astra, along with Grok 4.7, receive a recorded failure on criterion 4: “Applies a probe-dependence filter comparing probed channels (medium and heavy) against the unprobed light (DMSO background) channel to exclude non-specific binding.” Their reasons and interpretations differ, however: GPT-6 Astra explicitly questions the suitability of the background cutoff and keeps its conclusions exploratory, whereas Gemini 4 Argon uses stronger direct-target labels. A shared grading failure therefore does not imply identical scientific reasoning.

Meta Muse Wonder identifies PREPL and ACOT1, checks probe-background evidence, separates activity-associated capture from protein abundance, and proposes reporters with disrupted seed-matching sites. However, it acknowledges that testing separate fractions as independent replicates exaggerates the amount of independent information, then still uses significance categories from those tests in its combined candidate score. The warning is scientifically appropriate, but the ranking retains the problem the answer identified. Its interpretation is more cautious than a claim of direct validation, while the statistical support for the ranking still needs correction.

Grok 4.7 omits PREPL from its written shortlist, does not report the probe-background filter, and accepts some candidates supported by only 2 replicate blocks. Its selected response misses criterion 14: “Identifies PREPL as a top apparent direct target harboring an 8mer target site.” Here, “apparent” refers to a candidate suggested by the screening results, not a target whose direct regulation has been experimentally demonstrated. Missing PREPL changes which protein would be prioritized for follow-up. Grok 4.7 also calls ACOT1 and PPT2 the strongest direct targets without clearly stating that validation remains necessary.

All 5 model responses leave another comparison incomplete: whether the protein hits and computational sequence predictions agree more than expected by chance. All 5 miss criterion 12: “Statistically compares the proteomic hits to computational target predictions, such as through Fisher’s exact test or a rank-based comparison.” Identifying individual proteins that appear on both lists is useful, but it is a different question from testing whether the 2 analyses agree overall. A formal comparison would evaluate that agreement; it would still not replace an experiment testing direct regulation. The central biological distinction is between prioritizing a candidate and showing how it is regulated. A reporter experiment could compare the candidate’s normal regulatory sequence with a version in which the proposed miR-185 binding site is disrupted. Independent measurements of protein amount and enzyme activity would then help explain why probe capture decreased. Four models recover PREPL and ACOT1 as useful candidates, but they differ in replicate handling, background checks, statistical comparisons, and the strength of their claims. Those differences determine both which candidates are advanced and how confidently the researcher should interpret the shortlist.

### Plant Root Development Task Separating ORE15 Regulation from a Linked Genetic Change Task ID 673

This task asks why an Arabidopsis plant line carrying a disruption of ORE15 has shorter roots (Figure S6). Arabidopsis is a small flowering plant widely used in research. The altered line, ore15-2, has a smaller meristem, the region near the root tip where cells divide to support growth. It also shows changes in auxin and cytokinin responses, 2 hormone systems that regulate plant development. The models must explain these observations and select 15 genes for a follow-up experiment testing whether ORE15 associates with DNA near those genes. The supplied RNA-sequencing data measure RNA abundance in 3 root-tip samples from each plant type. Other observations include a weaker auxin-response reporter called DR5, less AUX1 protein, and a stronger cytokinin-response reporter called TCSn. A reporter is a designed readout of a biological response. The proposed follow-up, ChIP-qPCR, would test whether ORE15 is associated with DNA near selected genes. Choosing those genes requires considering whether their expression changes reflect ORE15 regulation or another difference carried by the altered plant line.

GPT-6 Astra, Gemini 4 Argon, Claude Opus 5.5, and Meta Muse Wonder recognize a block of neighboring genes on chromosome 1 whose RNA levels rise together. The genes have varied biological functions, yet their expression increases by roughly twofold. GPT-6 Astra reports 336 increased genes and only 1 decreased gene within this region. This spatial pattern is important: coordinated changes across neighboring genes can reflect a change in the chromosome segment itself, rather than ORE15 independently regulating every gene in the block. A linked duplication is one possible explanation, although confirming it requires measurements of genomic DNA. All 4 models exclude genes in this region from their proposed binding-test lists. That is a useful scientific decision because it avoids prioritizing genes whose increased expression may reflect the altered chromosome segment rather than direct regulation by ORE15. Grok 4.7 misses the neighboring expression pattern and includes genes from the region among its proposed direct-repression candidates. Its selected response misses criterion 8: “The response proposes 15 candidate genes for ORE15-GFP ChIP-qPCR of which none lie between AT1G24500 and AT1G31020.” The consequence is a different experimental shortlist, with some binding tests directed toward genes whose expression changes may have another cause.

Meta Muse Wonder identifies and excludes the block but leaves its connection to the ORE15 insertion site unexplained. Both Meta Muse Wonder and Grok 4.7 miss criterion 4: “The response states that the chromosome 1 region from about AT1G24500 to AT1G31020, which is raised in ore15-2, ends at the ORE15 insertion site, next to AT1G31040.” The location matters because it connects the unusual expression pattern to the region altered in this particular plant line. Meta Muse Wonder nevertheless proposes checking the DNA preserving a useful next step despite missing that specific connection. Grok 4.7’s omission is broader: it fails to use the block pattern to revise the candidate list.

Interpreting the hormone-related observations presents another difficulty in this. Gemini 4 Argon proposes that reduced expression of KMD genes allows ARR proteins to persist, strengthening cytokinin signaling. This offers a possible biological connection, but KMD RNA measurements alone do not show whether the relevant ARR proteins become more abundant or last longer. The RNA data also lack a consistent shift in the auxin-response and type-A ARR gene sets. GPT-6 Astra presents the KMD explanation as a hypothesis, but its discussion leaves out relevant alternatives, including the plant’s carbon status.

Gemini 4 Argon and GPT-6 Astra both miss criterion 13: “The response does not take the lower KMD genes as evidence of stronger cytokinin signalling and gives a reason, for example that KMD1 falls together with sugar-starvation genes such as ASN1, DIN10 and BT2, that type-A ARR genes show no consistent rise, or another valid biological or statistical argument.” Their answers differ in certainty, but neither fully addresses this requirement. The scientific issue is whether lower expression of 1 regulatory component is enough to explain the stronger cytokinin reporter particularly when other expression patterns offer a different interpretation.

Claude Opus 5.5 keeps several explanations open, including changes in where the response occurs within the root, changes at the protein level, and differences in nutrient state. Its selected response meets all recorded rubric requirements. This is useful reasoning because the hormone reporter, RNA measurements, and protein observations describe different aspects of the system. Meta Muse Wonder also considers protein production, movement, turnover, and cell composition, although it attributes some count-based sample differences to processing artifacts without independent technical information. Recognizing an unusual pattern is helpful; assigning its cause requires a test of the proposed explanation.

The next experiments should address these questions separately. Examining genomic DNA near ORE15 and comparing an independent mutant or a line with restored ORE15 function would help distinguish the intended gene disruption from a linked change in the original line. Genes outside the affected block provide a more informative starting list for binding experiments. Measurements of the relevant proteins, their distribution within the root, and the plant’s nutrient state would then help evaluate the hormone explanation. The main contrast is that 4 models use the neighboring expression pattern to improve the binding-test shortlist, while Grok 4.7 carries affected genes into that list. A separate difference concerns how confidently the models turn RNA changes into a hormone mechanism. The case shows why interpreting gene expression requires both biological knowledge and attention to where genes lie in the genome: the same increase in RNA can suggest regulation, altered gene copy number, or another feature of the experimental line, with different implications for the next experiment.

## Discussion

### Scientific Reasoning Connects Experimental Results to Research Decisions

BioResearchBench shows that even leading frontier AI models continue to struggle with challenging biological research questions. The highest overall success rate was 18%, yet the reviewed answers often contained useful calculations and biological insights. Models corrected sample assignments, accounted for delayed amino-acid labeling, and recognized DNA-cutting toxicity. However, they could complete important analytical steps and still misinterpret a measurement, overlook an alternative explanation, or propose an experiment based on an untested conclusion. In the protein-turnover task, a labeling complication was acknowledged but omitted from the main half-life calculation (Task ID 690). In the cytokine task, Meta Muse Wonder called the IL10 comparison provisionally not materially different within the samples that could be quantified and proposed a lower assay quantification limit. Because more treated than control samples could not be quantified, that subset did not resolve the comparison across all subjects (Task ID 732). These findings highlight the challenge of bringing data analysis and biological judgment together to guide research.

Biological measurements often provide only part of the information needed to explain a result. Oxygen consumption describes one aspect of cellular function, while membrane potential and cell abundance describe others. Protein capture can depend on enzyme activity as well as protein amount. RNA abundance, protein quantity, and ribosome occupancy reflect different processes. DNA sequencing can help predict which antibiotics a bacterium may resist, but a resistance-associated gene in a mixed sample may belong to another organism or be damaged and unable to function (Task ID 674). An incorrect resistance prediction could misdirect follow-up tests of which antibiotics inhibit the bacterium.

Several biological explanations can fit the same observation. A protein-labeling curve that differs from the measured cortical precursor can motivate investigation of an outside protein source, but it can also reflect a different precursor pool, mixed protein populations, or measurement error. A neighboring block of increased gene expression can suggest a linked genetic change rather than direct regulation of every gene in that region. Scientific judgment involves considering these alternatives and identifying which additional observation would most efficiently resolve the question. In the ovarian-cancer task (Task ID 701), Gemini 4 Argon interpreted lower HNF1B expression in bulk tissue as a change within the tumor cells, whereas GPT-6 Astra retained it as a bulk-marker difference. The measurement cannot separate fewer clear-cell epithelial cells in the sample from altered expression within those cells.

### Controls and Experimental Structure Determine What Can Be Concluded

In the CRISPR screen, 1 separately grown replicate culture was excluded because its guide-depletion measurements correlated less well with the other cultures and lacked their depletion pattern at regions with high DNA copy number. The remaining guides can still agree because cutting amplified DNA produces a shared growth defect, so guide agreement alone cannot distinguish DNA damage from loss of the targeted gene. In the mammary-cell sphere-formation task, a perturbation expected to increase sphere formation tests whether the assay can detect that response. It provides a check on assay responsiveness, while the specificity of the miR-424 effect requires separate testing.

Experimental structure also determines how much independent information an analysis contains. Nine LC–MS runs grouped into 3 replicate blocks are not equivalent to 9 independent biological replicates. Samples collected from different mice at different times produce a population time course, not a separate half-life measurement for every mouse. RNA reads assigned to the wrong sample can affect estimated changes even when the final biological classification happens to remain the same. These details connect statistical choices to the biological design. Recognizing them is insufficient if the subsequent calculation, ranking, or sample-size estimate ignores them. The proposed next experiment should address the unresolved explanation rather than assume it is correct. A perturbation that reduces gene function without cutting DNA helps investigate whether a CRISPR growth effect reflects the gene itself. An independent plant mutant helps test whether the phenotype follows the intended gene disruption rather than a linked change. A reporter with a disrupted microRNA-binding site tests whether regulation depends on that sequence. These are not interchangeable additions to an experimental plan: each is useful because its possible outcomes would change the interpretation of a specific result.

### Individual Rubric Failures Reveal More Than a Whole Task Score

Examining the specific requirements models missed was important for interpreting the benchmark. A whole-task failure records that an answer did not satisfy the grading procedure yet does not explain which parts remain useful or what needs correction. Models could recognize a filtering artifact in brain recordings or distinguish growth arrest from cell death while omitting requested analyses or producing inaccurate numerical estimates. Conversely, an answer could complete many calculations and still make a biological claim that changes the research decision. These observations argue for examining the content and consequence of a missed requirement and avoiding treating every failure as the same kind of error. The enzyme-kinetics task illustrates this distinction. GPT-6 Astra appropriately kept a preparation-selection decision conditional but omitted a requested numerical sensitivity calculation. Grok 4.7 supplied that calculation while distinguishing a multiplicative assay error from an additive background signal. Those assumptions have different consequences for the estimated kinetic parameters. The useful improvement was to show how the conclusion changed under an explicit assumption, rather than make a more confident claim from a single direct measurement. Measurements across substrate concentrations and checks of assay behavior remain necessary to characterize enzyme kinetics [18].

Rubrics make scientific requirements explicit, but they cannot capture every aspect of an answer. A criterion can combine several requirements, and a numerical expectation can depend on an analysis choice that the prompt permits the solver to make. In the cancer-screening task, a stricter guide threshold changed the number of genes classified as untestable. That difference affected the recorded grade, but the count discrepancy alone did not establish that the biological interpretation was wrong. The opposite also occurred: Claude Opus 5.5 and Gemini 4 Argon received full recorded criterion scores in the protein-turnover task while scientific review identified concerns about some interpretations of the fitted results. We retained the original grades and treated these observations as complementary scientific assessments rather than silently changing the scoring rule.

Evaluating model performance in biology requires choices about which analyses, explanations, and conclusions an answer should contain. During task development, experts were asked to explain how their reference answers could be verified and to provide supporting data, calculations, or sources, as appropriate. Authors and reviewers examined these materials alongside the questions and grading criteria through successive rounds of review. Even after repeated expert review, some choices remain open to reasonable disagreement. The task questions and grading criteria were explicit within the evaluation. Our aim was to understand how models approached biological questions, where their analyses were useful, and where their interpretations led to different conclusions or research decisions.

### Improvements Should Effect the Final Answer and Proposed Experiment

Several reviewed responses recognized a limitation but did not carry it through to the final output. An answer could acknowledge incomplete precursor labeling yet use a different model for its main half-life estimates. It could warn that measurements were not independent while retaining significance categories based on treating them as independent. It could report a residual functional deficit in a table while describing complete recovery in the conclusion. The half-life estimates need to incorporate precursor labeling, statistical support needs to respect the replicate blocks, and the recovery statement needs to retain the measured membrane-potential deficit.

Appropriate caution should also remain useful. Calling an amplified gene untestable can be reasonable when the screen cannot reliably separate gene loss from cutting damage. The answer should then identify an experiment that can address the question. Similarly, declining to assign a protein’s tissue of origin from an incompatible labeling curve is useful when accompanied by measurements that would test source alternatives.

Progress in biology and medicine can be slow and costly, often requiring repeated rounds of experimentation, analysis, and troubleshooting. When an experiment fails or produces unexpected results, researchers must decide which explanation to investigate and which next step is worth the time and resources. Ideally, frontier AI models would help prioritize the analyses and experiments most likely to answer the scientific question, reducing time and materials spent on less informative approaches. The aim is to explain the scientific meaning of the results and identify the next most useful analysis or experiment for the researcher.

Planning experiments requires the same discipline. A difference between mouse genotypes can be used as an explicitly stated planning scenario, but it is not a measured treatment effect. An animal study needs an outcome that can be measured in individual animals, a treatment effect worth detecting, and an estimate of between-animal variability. Pilot measurements can inform feasibility and variability, while sensitivity analyses show how the required sample size changes across assumptions [19]. Returning a sample-size number without making those choices explicit can obscure the most important biological uncertainty.

The differences among model responses suggest opportunities for targeted checking. One model may supply a calculation another omitted or preserve a biological qualification another lost. However, agreement among models is not independent validation: all 5 selected responses made the same whole-field exclusion after a nuclear-segmentation problem in the respiration task. Any benefit from combining answers would need to be tested against the measurements and task requirements. We did not evaluate a multi-model workflow in this study.

These findings motivate training and evaluation that emphasize the connection between analysis and decision. Expert examples could show how researchers weigh conflicting measurements, recognize alternative explanations, choose informative controls, and revise a conclusion when assumptions fail. Such examples should explain why a decision follows from the experimental details, including situations in which more than 1 next step is reasonable. Whether a particular training approach improves these behaviors remains to be tested.

### Relevance to developing AI research assistants and use of frontier AI in science

For model developers and research laboratories, these findings suggest concrete behaviors to evaluate in systems used for scientific research. Future evaluations should examine how researchers and models work together over several interactions or “turns” with a model. Researchers may challenge an interpretation, provide additional results, or refine the question as the work develops. Evaluations could assess whether the model responds appropriately to that input, corrects mistakes, and helps the researcher choose a useful next step. Future evaluations could also follow models through sustained research projects with experimentally testable goals. Examples include developing an antibody or protein binder with a specified function, identifying an antimicrobial candidate, or determining the correct culture conditions for a microbe that is finicky to grow. Researchers and models would work through successive rounds of analysis, experimentation, and troubleshooting, with assessment tied to the results obtained and the time and resources used. Comparisons with researchers working without AI would help determine where the model made a useful contribution. Project-based evaluations could complement task collections by examining whether models help turn an initial scientific question into reproducible progress.

A separate priority is to develop ways of assessing open-ended scientific work beyond fixed rubrics since the reality of the most challenging biology and medicine tasks are that they may be open to interpretation with multiple correct answers. These could include independently reproducing an analysis, testing predictions against new data, experimentally evaluating proposed explanations, and asking experts to compare the usefulness of different responses. Such approaches could recognize sound analyses and ideas that were not anticipated when the grading criteria were written.

We recognize the scientific and technical work involved in developing biological benchmarks including LABBench2 from Edison Scientific [8], LifeSciBench from OpenAI and Tacit Labs [9], OmicsBench from LatchBio [10], BioMysteryBench from Anthropic [13], CompBioBench from Genentech and Roche [20], and AstaBench from the Allen Institute for AI [21], among many others. We also recognize the Stanford led Terminal Bench Science collaboration [14] for bringing together domain experts to develop verifiable evaluations of scientific research workflows. These contributions and the continued exchange of ideas help the research community develop more informative evaluations of frontier AI models in biology and medicine. Advancing frontier AI models in biology and medicine is a collective effort involving data companies, AI developers, academic researchers, and scientific organizations. Developing useful evaluations draws on their combined expertise in biology, data, and model development.

## Conclusion

Across the tasks examined in detail, models often completed useful analyses but differed in how they interpreted the results and answered the biological question. BioResearchBench documents both useful scientific contributions and consequential errors in frontier AI responses to realistic biological research questions. Low whole task success did not erase the value of corrected sample labels, appropriate labeling models, recognized screening artifacts, or informative biological classifications. At the same time, those successes did not guarantee that the final explanation or proposed experiment was appropriate.

Other frontier AI model conclusions went beyond the measurements and data provided by the task. A poor fit to cortical labeling was treated as proof of an outside protein source, and decreased enzyme capture together with a sequence match was treated as direct microRNA regulation. In the respiration case, 1 recovery statement contradicted a deficit reported in the answer’s own table. These errors need different corrections: a narrower interpretation, a test that distinguishes competing explanations, or agreement between the summary and the reported result. The follow-up experiment should address the specific unresolved cause. Reducing a gene’s function without cutting DNA helps test whether a CRISPR growth effect reflects the gene or cutting injury. Comparing an independent plant mutant helps separate ORE15 disruption from a linked change. Disrupting a proposed microRNA-binding site tests whether the response depends on that sequence. We also conclude that different models are better at different capabilities and it may be worthwhile for scientists to use multiple frontier AI models in their research.

Collections such as BioResearchBench offer a focused view of how models approach biological research: how they use scientific tools and databases, perform calculations and statistical analyses, work with imperfect data, and decide what to do next. The tasks also ask models to bring these steps together to explain what the results mean and draw conclusions supported by what was measured.

Examining their responses in detail reveals useful capabilities alongside specific gaps in interpretation and scientific judgment. Task-based benchmarks can help identify these gaps, while evaluations of sustained research projects can test how models contribute as questions evolve and new results arrive. Together, these approaches can guide the development of AI systems that are more useful to researchers in biology and medicine.

### Limitations of this work

The collection was deliberately selected for difficulty. Task acceptance required a failed GPT-6 Astra response during development and some questions were revised or removed as newer models answered them correctly. We selected GPT-6 Astra at the time since it was one of the strongest biology models and we wanted to use a strong model for our threshold in difficulty. The reported rates therefore characterize this selected collection rather than an unbiased sample of biological research questions or a general ranking of model capabilities. The final 50 were selected following review of expert-developed tasks for scientific accuracy, clarity, difficulty, and biological and skill coverage, rather than ranking tasks by model success rates.

Reviewers sometimes made questions more explicit about assumptions, required comparisons, and expected outputs so that answers could be judged consistently. This reduced avoidable ambiguity, but made some questions more prescriptive than those a biologist would normally pose during research. The tasks therefore test performance on carefully specified research problems, not the full process of deciding how an initially open-ended problem should be framed. Grading should accommodate scientifically justified alternative approaches that satisfy the task’s requirements.

## Acknowledgements

We are deeply grateful to the biology experts whose knowledge and research experience made BioResearchBench possible. They developed realistic research questions, prepared supporting materials and reference answers, and helped refine the tasks and grading rubrics through detailed scientific review.

We sincerely thank the biology domain leads, Sabah Ul-Hasan and Stelios Lamprou, for their scientific guidance, contributions to task review, and support for the experts throughout this project.

## Data Availability

The full 50-question benchmark is available for commercial licensing through Office Hours. Requests for sample tasks or access to the benchmark can be submitted through the <u>Office Hours Live</u> <u>Benchmark Results Page</u> or by emailing.

## Competing Interests

Several authors are employed by Office Hours which offers commercial licensing of BioResearchBench. The authors declare no other competing interests.

## Use of AI

This manuscript was developed under human direction with AI assistance in analysis, review, and editing. AI assistance in rubric development and automated response grading is described in Materials and Methods. AI was also used to create some figures from human-provided data with deep expert guidance.

## BioResearchBench Supplementary Figures

**Figure S1.**
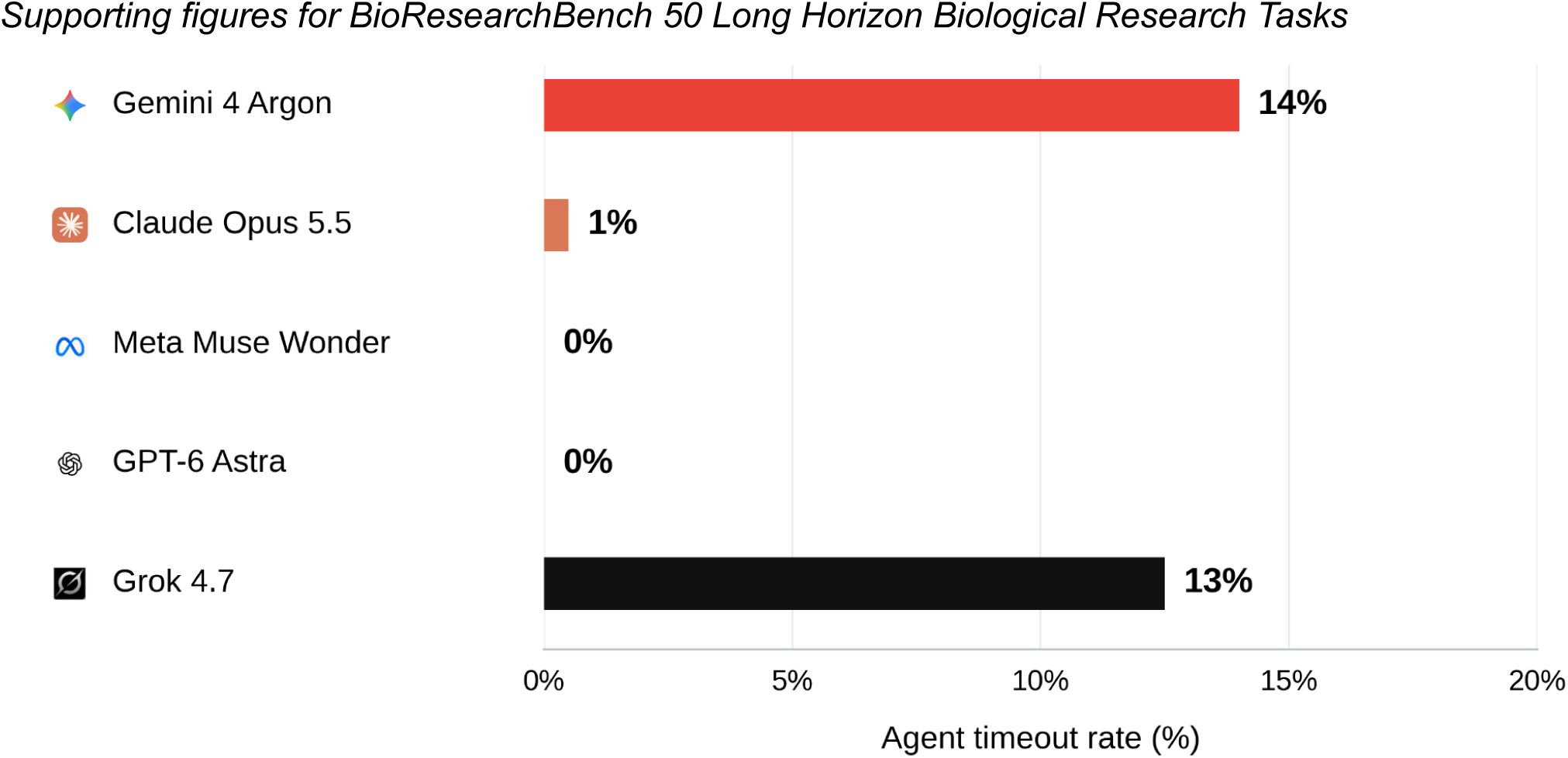
Agent timeout rate. Agent timeouts occurred in 14% of Gemini 4 Argon trials, 1% of Claude Opus 5.5 trials, 0% of Meta Muse Wonder trials, and 13% of Grok 4.7 trials: 28, 1, 0, and 25 of 200 assigned trials, respectively. No agent timeouts were reported for GPT-6 Astra (0%). Timeouts are reported separately from scientific grading failures and other operational errors. Bar lengths use exact fractions and labels use whole percentages. **Significance:** Agent timeouts were more frequent for Gemini 4 Argon and Grok 4.7 than for Claude Opus 5.5, Meta Muse Wonder, and GPT-6 Astra under the tested configurations. A timeout can interrupt an analysis before a finished answer is available for scientific evaluation. Timed-out attempts were excluded from task success calculations to be fair and to actually test biology answers that models gave as correct or wrong.

**Figure S2.**
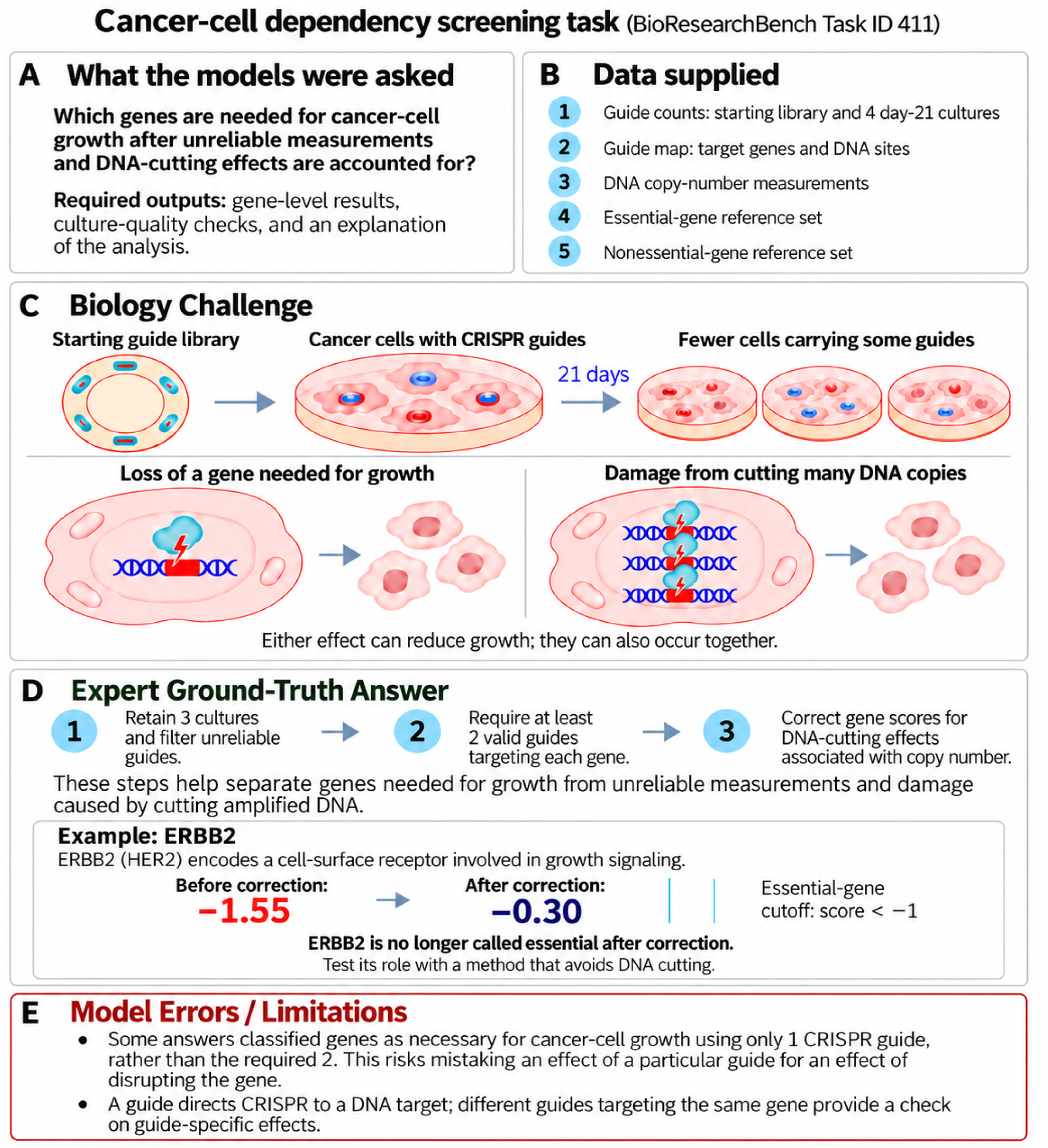
Cancer cell dependency screening task Task ID 411 Illustration. Models were asked to identify genes needed for cancer-cell growth using CRISPR screening data. Guide RNAs direct Cas9 to DNA targets, and sequencing measures each guide’s representation before and after 21 days of growth. Cells carrying a guide may become less abundant because disrupting its target gene impairs growth or because cutting many copies of an amplified DNA region causes damage. The expert ground-truth answer combines culture-quality checks, guide filtering, and copy-number correction before classifying genes. ERBB2 encodes HER2, a cell-surface receptor involved in growth signaling. Its score changes from −1.55 to −0.30 after correction, moving above the task’s essential-gene cutoff of −1. Testing its role without DNA cutting would help distinguish gene function from cutting-related damage. Panel E describes a limitation found in a model response: some gene classifications used only 1 retained guide rather than the required 2, leaving them more vulnerable to guide-specific effects.

**Figure S3.**
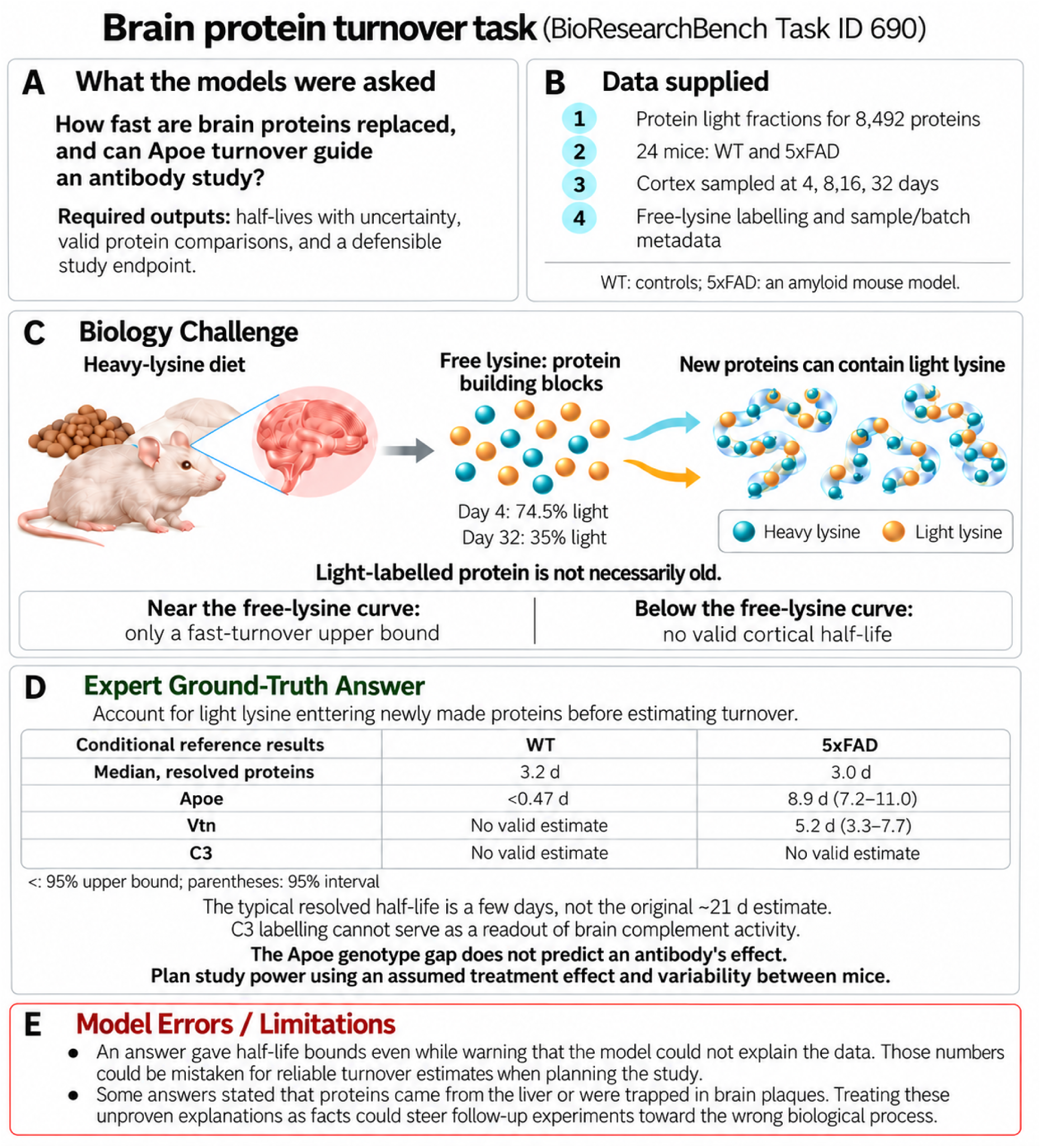
Brain protein turnover task Task ID 690 Illustration. Models were asked to estimate how quickly proteins are replaced in mouse cortex and assess Apoe turnover as a possible endpoint for an antibody study. Wild-type controls and 5xFAD mice, which develop amyloid pathology, received heavy-isotope-labelled lysine and were sampled at 4 time points. Lysine is a building block of proteins. Because the available lysine remained partly unlabelled, newly synthesized proteins could also remain partly unlabelled. The expert ground-truth answer accounts for this when estimating half-lives. Under its single-pool model, proteins closely tracking the precursor can support only an upper bound on half-life, while curves significantly below the precursor are incompatible with that model. Panel D shows selected conditional reference estimates; parentheses denote 95% intervals and “<” denotes a 95% upper bound. The Apoe difference between genotypes cannot by itself specify an antibody’s treatment effect. Panel E describes numerical and mechanistic claims found in selected responses that could mislead study planning.

**Figure S4.**
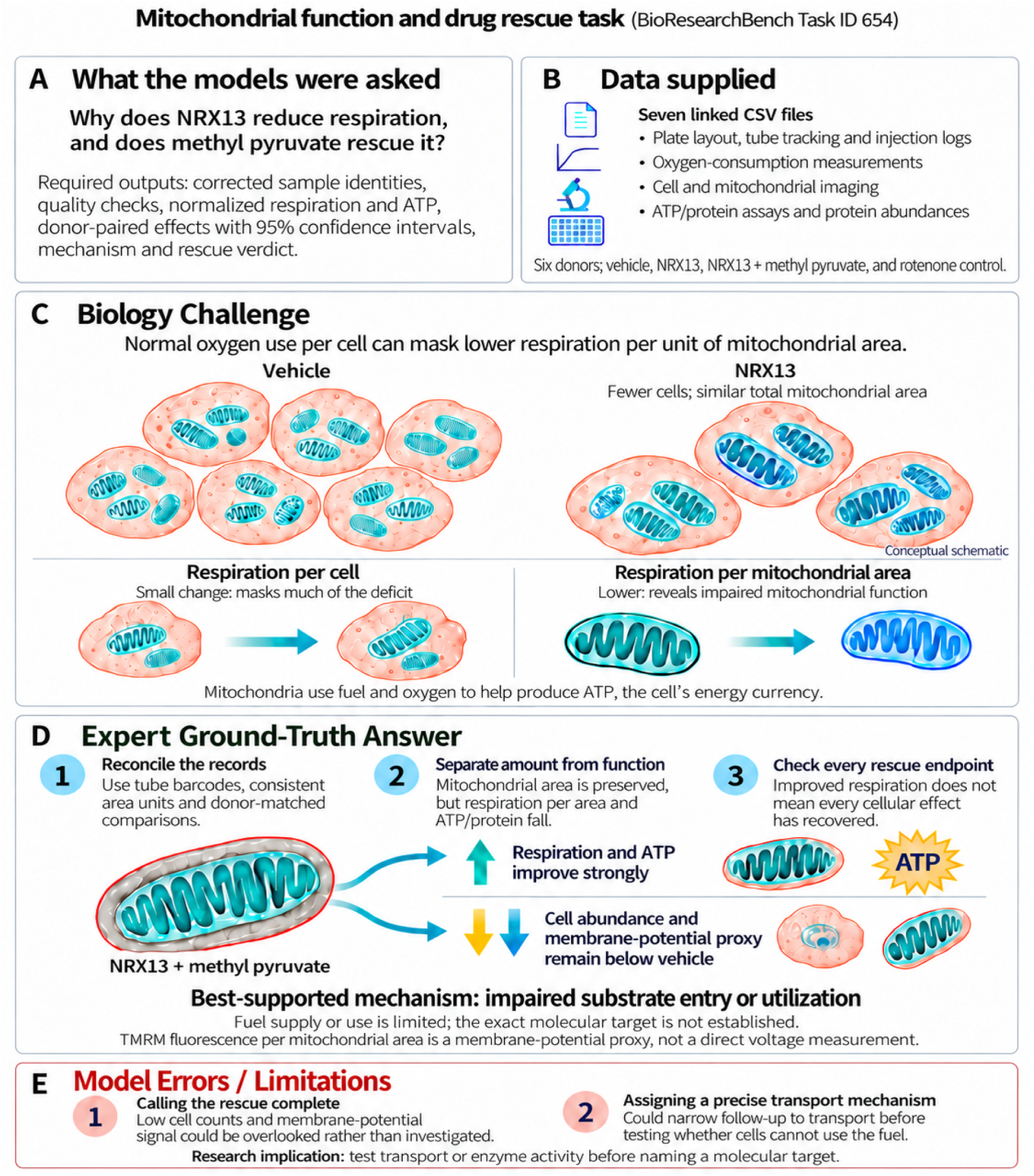
Mitochondrial function and drug rescue task Task ID 654 Illustration. Models were asked why NRX13 reduces respiration and whether methyl pyruvate reverses its effects. Seven linked data files provide measurements from 6 donor-matched preparations, including oxygen consumption, cell and mitochondrial imaging, ATP, and protein abundance. The expert ground-truth answer reconciles sample identities, imaging units, and quality checks before comparing conditions. NRX13 reduces cell abundance while total mitochondrial area remains broadly preserved. Dividing respiration by cell count can therefore hide a larger loss of function per mitochondrial area. Methyl pyruvate strongly improves respiration and ATP, but cell abundance and the membrane-potential measurement remain below the vehicle control. The results favor impaired fuel entry or use without identifying an exact molecular target. Panel E shows how claims of complete rescue or a specific transport mechanism could lead researchers to overlook remaining effects or narrow follow-up experiments prematurely. The cell diagrams are conceptual, not quantitative counts.

**Figure S5.**
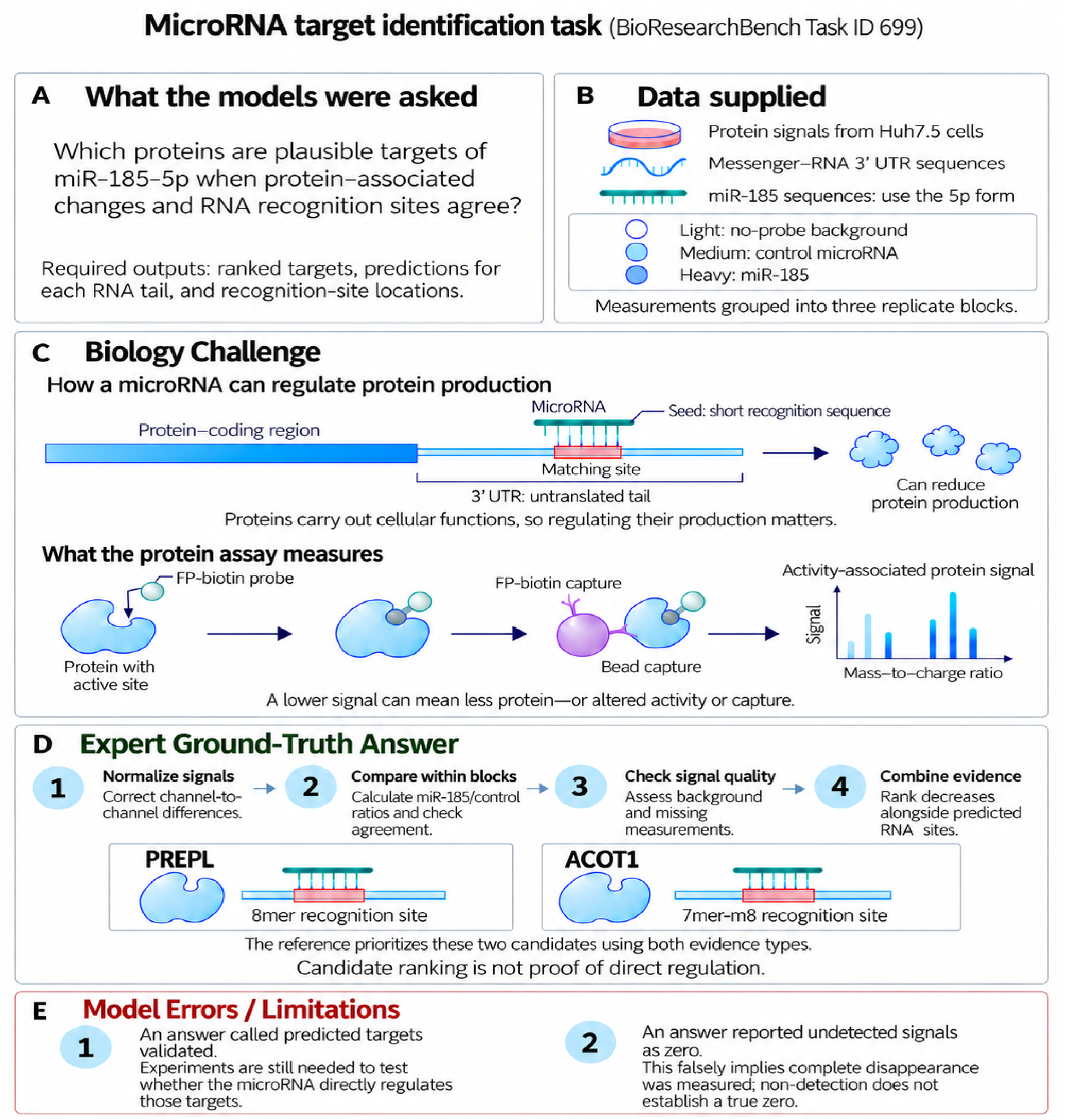
MicroRNA target identification task Task ID 699 Illustration. Models were asked to combine protein measurements from Huh7.5 cells with predicted recognition sites for miR-185-5p and produce ranked targets, transcript predictions, and site locations. MicroRNAs can reduce protein production by recognizing complementary sequences in messenger RNA, often within the 3′ untranslated region, a tail that is not translated into protein. The FP-biotin probe captures proteins according to their activity and accessibility to the probe, so the resulting signal also depends on factors beyond protein amount. The light channel measures no-probe background; medium and heavy represent control-microRNA and miR-185 conditions. Nine runs are grouped into 3 replicate blocks. The expert ground-truth answer normalizes signals, checks background and missing measurements, and combines reproducible decreases with sequence matches to prioritize PREPL and ACOT1. Experiments would still be needed to test direct regulation. Panel E shows 2 errors found in a selected response: describing predicted targets as validated and representing undetected signals as zero.

**Figure S6.**
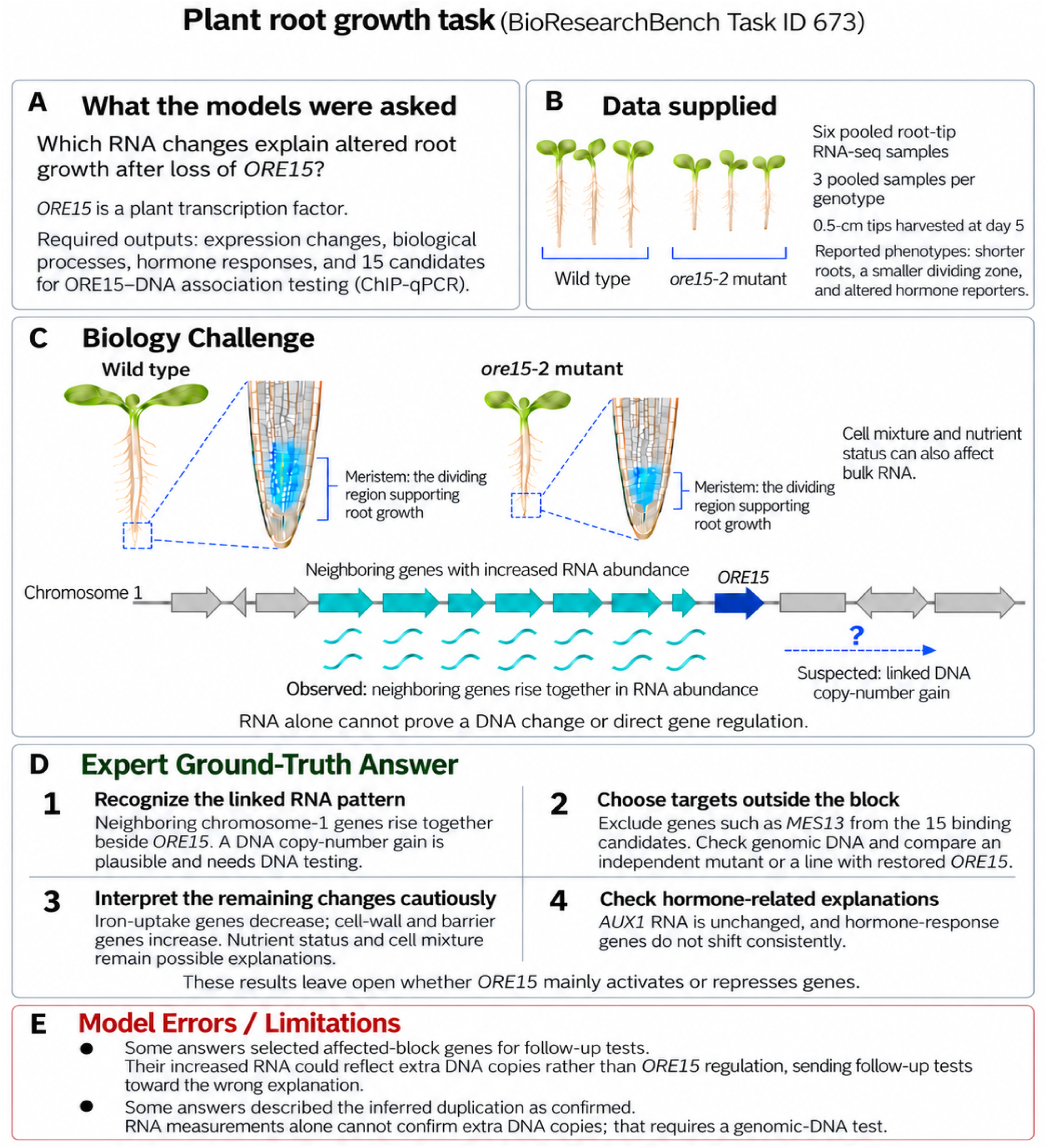
Plant root growth task Task ID 673 Illustration. Models were asked to interpret RNA measurements from Arabidopsis root tips after loss of ORE15, a protein that regulates gene expression, and select 15 candidate genes for follow-up binding tests. The mutant plants had shorter roots and a smaller meristem, the region where cells divide to support root growth. The expert ground-truth answer identifies a block of neighboring chromosome-1 genes whose RNA levels rise together. Extra copies of that DNA region could explain the pattern, so increased RNA alone should not be attributed to ORE15 regulation. The answer recommends testing genomic DNA, selecting binding candidates outside the affected block, and comparing an independent mutant or a line with restored ORE15. Changes in nutrient status and the proportions of different cell types could also contribute to the remaining RNA differences. Panel E describes selected responses that overlooked the affected block when choosing follow-up targets or reported the suspected duplication as confirmed.

## Notes

https://horizon.officehours.com/research/a-50-question-biology-benchmark-2026-10-21

### Competing Interest Statement

The authors have declared no competing interest.

